# Developmental and age-related synapse elimination is mediated by glial Croquemort

**DOI:** 10.1101/2024.06.24.600214

**Authors:** Taylor R Jay, Yunsik Kang, Victor Ouellet-Massicotte, Mariel Kristine B Micael, Victoria L Kacouros-Perkins, Jiakun Chen, Amy Sheehan, Marc R Freeman

## Abstract

Neurons and glia work together to dynamically regulate neural circuit assembly and maintenance. In this study, we show *Drosophila* exhibit large-scale synapse formation and elimination as part of normal CNS circuit maturation, and that glia use conserved molecules to regulate these processes. Using a high throughput ELISA-based *in vivo* screening assay, we identify new glial genes that regulate synapse numbers in *Drosophila in vivo*, including the scavenger receptor ortholog Croquemort (Crq). Crq acts as an essential regulator of glial-dependent synapse elimination during development, with glial Crq loss leading to excess CNS synapses and progressive seizure susceptibility in adults. Loss of Crq in glia also prevents age-related synaptic, but not neuronal loss, in the adult brain. This work provides new insights into the cellular and molecular mechanisms that underlie synapse development and maintenance across the lifespan, and identifies glial Crq as a key regulator of these processes.

## Introduction

Establishing and maintaining appropriate synaptic connections is critical for nervous system function. During development, synapse formation and selective synapse elimination act together to construct this precise synaptic circuitry. Disruption in the process of synapse formation has been implicated in intellectual disability^1^, injury-induced pain^2^, and failure to recover after stroke^3^. Disruption in synapse elimination is associated with epilepsy^4^, autism^5^ and schizophrenia^6^, and aberrant elimination of synapses has been proposed to lead to cognitive decline with aging^7^ and in neurodegenerative diseases^8^. Understanding mechanisms that contribute to the formation and elimination of synapses will therefore be critical to understand normal development and neurological disease.

Over the past two decades, we have come to appreciate that neurons do not orchestrate synapse development alone. Glia are essential regulators of both synapse formation and elimination in development and disease^9,10^. For example, glial *TSP1/2*^11^ and *SPARC*^12^ were found to be required for excitatory synapse formation and maturation, and *MEGF10*^13^ and complement^14,15^ are necessary for synapse elimination and circuit refinement. Many of these molecules were initially found by growing subsets of neurons *in vitro*, with or without glia conditioned media, to identify secreted glial factors that regulate synapse development^16^. Others were found using targeted approaches, identifying candidates via screens in model organisms, transcriptional profiling or disease association studies, and then validating the role for these genes in synapse development using mouse models^13,15,17^.

Biochemical studies in mammals have allowed us to learn primarily about the secreted molecules that glia employ to regulate development of synapses in excitatory neurons. However, contact-dependent mechanisms also play a role, though the molecular basis remains poorly defined^18,19^. Recent work demonstrates that glia also regulate inhibitory synapse development, though how glia regulate non-excitatory synapses remains relatively understudied^20,21^. Finally, it is important to note that *in vitro* systems cannot recapitulate endogenous patterns of cellular interactions, neuronal activity and biological age. We are therefore far from having a complete picture of neuron-glia signaling during establishment of circuit connectivity, and many *in vivo* synaptic regulatory molecules await discovery.

Addressing these questions will require large-scale gene discovery efforts in an *in vivo* system, and manipulation of gene expression in glia or neurons while reading out the effect on synapses. In this study, we use *Drosophila* for such a functional screen and provide evidence that *Drosophila* exhibit continued synapse formation and selective elimination throughout early adulthood, akin to the processes that occur during circuit maturation in mammals, as well as age-related synaptic loss. We identify 48 new genes that are required in glia *in vivo* for proper synapse development, including Crq, a scavenger receptor ortholog and part of an immune receptor family not previously implicated in synapse development. Loss of Crq in glia was sufficient to suppress glial synapse engulfment, resulting in increased synapse number and ultimately seizure susceptibility. Intriguingly, loss of Crq was sufficient to prevent age-related synaptic loss, in contrast to the well-studied engulfment receptor Draper (Drpr) / MEGF10, arguing for selective use of glial engulfment receptors during synapse loss in the aging brain.

## Results

### *Drosophila* use conserved glial mechanisms to regulate synapse development and maintenance across the lifespan

Formation of synapses that comprise the circuitry of adult *Drosophila* begins at late pupal stages^22^. We defined the dynamics of synaptic changes in *Drosophila* across their adult lifespan, and determine whether glia regulate synapse numbers (Fig 1A). As a proxy for measuring synapses, we evaluated levels of Brunchpilot (Brp), a molecule present in presynaptic active zones^23^, and the canonical synaptic marker in the fly. We used an existing fly line in which Brp is endogenously tagged with GFP (Fig 1B) and measured GFP concentration in whole adult heads by ELISA^24^. Based on this assay, we found that, like mammals, adult flies appear to undergo defined periods of synapse formation and elimination during early adulthood (Fig 1C), with increases in Brp-GFP levels evident between eclosion (0 days post-eclosion – dpe) and early adulthood at 3 dpe, and a decrease observed between 3 dpe and mid-adulthood (10-12 dpe). Measurement of additional pre- (Fig S1A) and post-synaptic (Fig S1B) markers also showed a decrease in synaptic protein levels after eclosion. *Drosophila* also exhibit synaptic loss in aged 30-35 dpe flies (Fig 1C), akin to the synaptic loss observed with age in mammals^25-28^. Thus, flies display very similar dynamics in synaptic marker levels across the lifespan as mice and humans.

**Figure 1:**
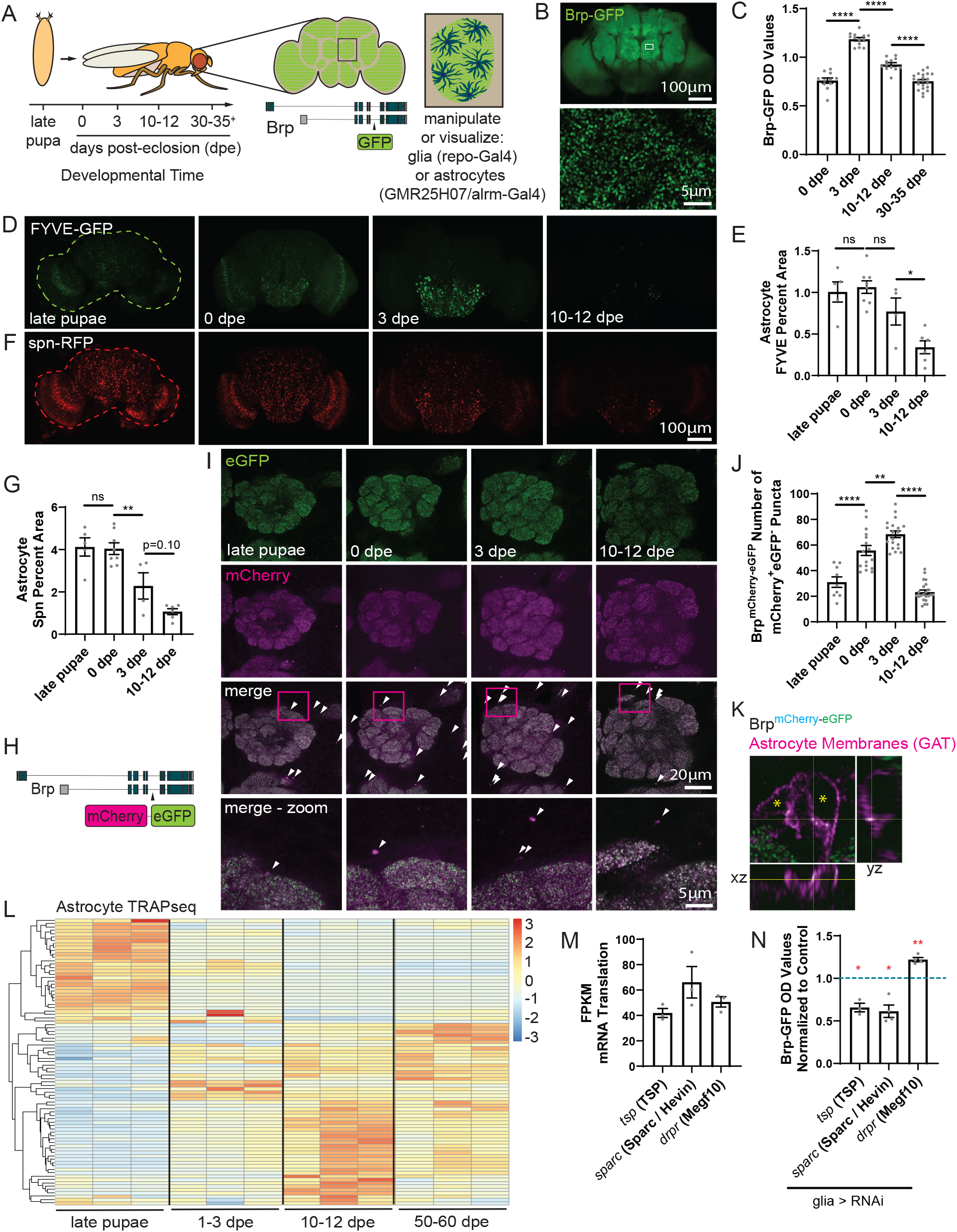
*Drosophila* use conserved glial mechanisms to regulate synapse development and maintenance across the lifespan. (A) Formation of the adult fly circuitry begins at late pupal stages (48-96 hr after puparium formation) and in this study we evaluated glial and synaptic changes that occurred from this time point through the rest of the fly lifespan (30-35 days post-eclosion (dpe)+). To visualize and evaluate synapses, the synaptic protein Brp was endogenously labeled with GFP (zoomed out image acquired with a 10x objective and inset acquired with a 63x objective) (B), and gene expression was manipulated in all glia using *repo-Gal4* or just in astrocytes using *GMR25H07-Gal4* or *alrm-Gal4*. (C) Brp-GFP protein levels were analyzed by ELISA in fly heads (*brp-GFP/+, repo-Gal4/+*; males and females combined) collected at the indicated time points. For each sample, 3 fly heads were pooled, and each n represents the optical density (OD) value for that sample. The n for each group are: 11 (0 dpe), 13 (3 dpe), 12 (10-12 dpe), 21 (30-35 dpe). Data from all groups are normally distributed and compared using an ordinary one-way ANOVA (****) with Sidak’s post-hoc comparisons indicated. (D) Images were acquired using a 10x objective, with 12 z sections, beginning at the anterior-most part of the brain and taken 5μm apart from fly brains in which astrocytes express FYVE-GFP, a PI3P marker, which labels endosomes and early phagosomes (*alrm-Gal4/+, UAS-FYVE-GFP/+, UAS-spn-RFP*, males and females combined). (E) Images were z projected through the entire set of z planes acquired, thresholded and the percent FYVE+ area measured across the entire brain area (outlined with a green dotted line). These values are reported as raw percentages and are not normalized to any group. The n for each group are: 5 (late pupae), 8 (0 dpe), 4 (3 dpe), 6 (10-12 dpe). All groups were normally distributed and compared using an ordinary one-way ANOVA (***) with Sidak’s post-hoc comparisons indicated. (F) The late phagosomal marker spn-RFP was expressed in astrocytes (*alrm-Gal4/+, UAS-FYVE-GFP/+, UAS-spn-RFP/+*, males and females combined) and imaged and processed as described for FYVE above. (G) The percent spn^+^ area was measured across the entire brain (outlined with a red dotted line). These values are reported as raw percentages and are not normalized to any group. The n for each group are: 5 (late pupae), 8 (0 dpe), 4 (3 dpe), 6 (10-12dpe). All groups are normally distributed and compared using an ordinary one-way ANOVA (****) with Sidak’s post-hoc comparisons indicated. (H) We developed a tool to evaluate synaptic engulfment in which a pH-insensitive mCherry and a pH-sensitive eGFP were knocked into the endogenous Brp locus. (I) eGFP and mCherry signal were visualized across the entire central brain area of *brp*^*mCherry-eGFP*^ / + flies (males and females combined). Brains were imaged with a 40x objective and z planes were acquired starting at the anterior-most portion of the brain and taken every 5μm for 8-10 z planes. (J) The number of mCherry^+^eGFP^-^ puncta were quantified across the central brain in each z section and the average number of mCherry+eGFP-puncta / section was calculated. Each n represents this value from one individual brain. The n for each group are: 9 (late pupae), 16 (0 dpe), 19 (3 dpe), 22 (10-12 dpe). All groups were normally distributed and compared using an ordinary one-way ANOVA (****) with Sidak’s post-hoc comparisons indicated. (K) Flies expressing Brp^mCherry-eGFP^ (brp^mCherry-eGFP^ / +) were collected at 3 dpe and brains were stained for the astrocyte membrane marker GAT. Images were acquired using a 63x objective with optical sections of 0.2μm. Localization of mCherry+eGFP-puncta within astrocytes was assessed. (L) We performed TRAP sequencing to evaluate genes translated in astrocytes across late pupal stages (48-72hr APF) through the adult lifespan using male and female *alrm-Gal4/+, elav-Gal80/ +, UAS-L10a-eGFP/+* flies. Differences in expression across the 100 most differentially expressed genes (red - high, blue - low) was plotted. (M) FPKM (Fragments Per Kilobase of transcript per Million mapped reads) values were extracted from the TRAPseq dataset for tsp, sparc and drpr. Mammalian orthologs are listed in parentheses in the x-axis labels. (N) RNAi’s were used to knockdown each of these genes in glia and Brp-GFP levels were measured by ELISA. Three fly heads were pooled for each sample from *brp-GFP/+, repo-Gal4/+, UAS-RNAi/+* flies, along with controls (*brp-GFP/+, repo-Gal4/+*). OD values from each experimental sample was normalized to the control average. The n for each group are: 3 (tsp), 4 (sparc) and 4 (drpr). All groups were normally distributed and each experimental condition was compared using a one sample t-test relative to the hypothetical value of 1.

We next sought to determine whether glia play a role in regulating these dynamic changes in synapses across the *Drosophila* lifespan. Glia have been previously implicated in regulating synapse formation in the fly, as partial ablation of astrocytes during pupal development resulted in reduced synapse numbers in adults^22^. To assess whether glia might also regulate developmental synapse elimination, we evaluated whether and when glia engage in phagocytic activity across late pupal through adult stages. Using genetically encoded markers in astrocytes, we observed FYVE-GFP, a PI3P marker which labels endosomes and has previously been used to monitor engulfment in the fly^29^ (Fig 1D, E) and spn-RFP, a phagolysosomal marker across the entire fly brain (outlined by dashed lines) (Fig 1F, G). This signal was present broadly across the brain at late pupal stages, and was maintained within 1 day of eclosion (0 dpe). These phagocytic markers decreased substantially by 10-12 dpe (for FYVE, a 6% increase from late pupal stages to 0 dpe, a 28% decrease between late pupal stages and 3 dpe and a 56% decrease between late pupal stages and 10-12 dpe; for spn, a 2% decrease from late pupal stages to 0 dpe, a 43% decrease between late pupal stages and 3 dpe and a 53% decrease between late pupal stages and 10-12 dpe). These data are consistent with the notion that glia are actively phagocytic at times when synapse elimination occurs.

To more directly monitor synaptic engulfment, we adapted a transgene which had previously been used to evaluate synaptic engulfment in mammalian astrocytes^30^ and generated a fly in which Brp was endogenously dual-labeled with a pH-insensitive mCherry and a pH-sensitive eGFP (Fig 1H). Using this tool, synapses appear dual labeled until they are engulfed and trafficked to acidic lysosomes, where the eGFP signal was quenched and only mCherry fluorescence remains. We validated this tool by showing it produced expected changes in eGFP (Fig S2A, B) and the ratio of eGFP and mCherry fluorescence (Fig S2A, C) during metamorphosis, when glia prodigiously engulf synaptic material. We quantified the number of mCherry^+^ eGFP- puncta from late pupal through early adult stages (Fig 1I). Engulfed synaptic material was present at late pupal stages, and this increased over the course of early adulthood, peaking at 3 dpe before dropping dramatically at 10-12 dpe (Fig 1J, Fig S2D-F). We were able to visualize both lysosomes (Fig S2G) and mCherry+eGFP-puncta within astrocyte membranes (Fig 1K, Fig S2H). Together, these data support the notion that *Drosophila* glia engulf synapses during early adulthood, coincident with a period of synapse elimination.

We next evaluated whether fly glia use similar molecular pathways to regulate synapse numbers in comparison to astrocytes in mammals. We first analyzed a TRAPseq dataset we generated^31^ to assess changes in astrocyte gene translation across the developmental time points we identified as important periods of synapse formation and elimination (Fig S3A, Fig 1L). We found that orthologs to genes associated with synapse formation, tsp (*TSP1/2*)^11^ and *sparc* (*Sparc / Hevin*)^12^, and synapse elimination, *drpr* (*Megf10*)^13^, were expressed in astrocytes during the developmental times when these processes occur in the fly (Fig 1M). We knocked each of these genes down in glia and measured the effects on levels of the synaptic protein Brp and found glial knockdown of *tsp* and *sparc* resulted in a reduction in synaptic protein levels, while knockdown of *drpr* resulted in an increase (Fig 1N).

While the strongest changes in the astrocyte translatome were present between late pupal stages and after eclosion (Fig S3B), there were still many genes that were differentially translated in astrocytes at early adulthood (1-3 dpe) compared to those at mid-adulthood (10-12 dpe) (Fig S3C). We also identified changes in gene expression in astrocytes after aging animals 50-60 days (Fig S3D, also see Table S1). This dataset provides a rich database to define mechanisms underlying changes in astrocyte phenotype across the lifespan. Moreover, our data argue that *Drosophila* glia exhibit conserved expression and function of previously established mammalian glial regulators of synapse development, and flies should therefore by a useful system in which to identify new glial regulators of synapse numbers.

### An *in vivo* genetic screen identifies glial regulators of synapse development

We used available transcriptional and proteomic datasets to identify genes and proteins expressed in mammalian glia,^32-47^ identified the fly orthologs of those genes, and screened them for roles in synapse formation or elimination *in vivo*. For each candidate, identified up to two RNAi lines available in public repositories^48^ that would allow us to knockdown those genes, resulting in 695 genes and 971 screening lines (Table S2).

We took advantage of a screening paradigm we recently developed, using ELISA-based measurement of GFP levels in fly lysates to quantitatively measure levels of endogenously GFP-tagged proteins^24^. Specifically, we used this ELISA method to measure levels of the endogenously tagged synaptic protein, Brp-GFP as a proxy to evaluate synapses (Fig 2A). We found that we could reliably measure Brp-GFP signal from as little as ¼ of a fly head (Fig S4A, B). To minimize variability, we used male flies (Fig 42C), pooled 3 fly heads per sample and ran 3 biological replicates per line (Fig S4D). It had been previously reported that there are circadian changes in synapse levels in flies, as measured by Brp protein levels^49^. We found similar results for flies at 10-12 dpe and these changes were abrogated in *per*^01^ mutants in which circadian rhythms are disrupted^50^ (Fig S4E). We did not observe significant circadian changes in synapses earlier in development, at 3 dpe (Fig S4F). Therefore, we chose to perform the screen at 3 dpe as it appeared to be an ideal time point to capture genes that affected dynamics of both synapse formation and elimination, and to avoid the potential confounding effect of circadian synaptic changes.

**Figure 2:**
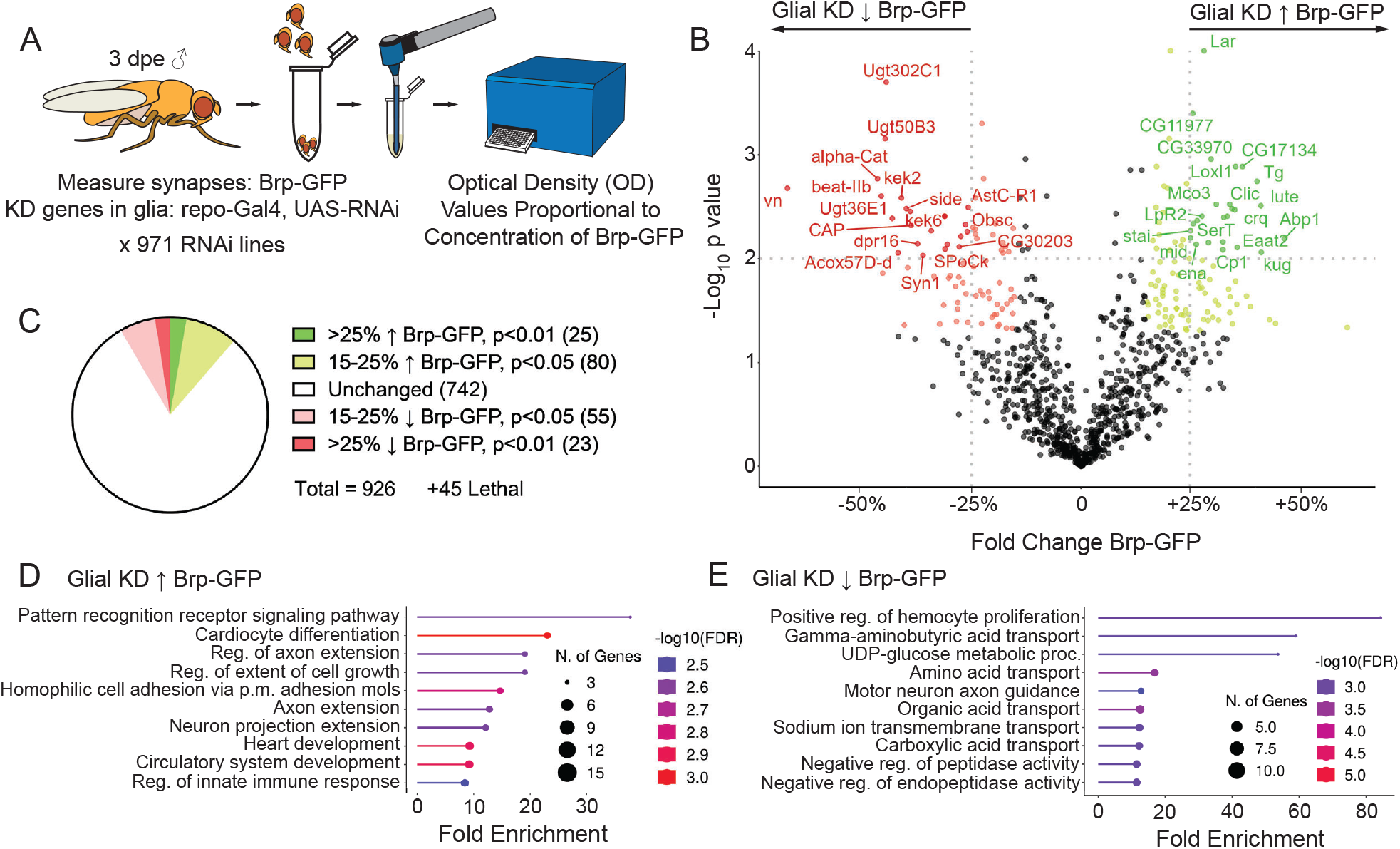
An *in vivo* genetic screen identifies glial regulators of synapse development. (A) A screen to identify glial regulators of synapse development was performed in male *Drosophila* at 3 dpe. Flies expressing Brp-GFP and the glial driver *repo-Gal4* were crossed with 971 selected UAS-RNAi lines (generating experimental flies with the genotype *brp-GFP / +, repo-Gal4 / +, UAS-RNAi / +)* For controls, *brp-GFP, repo-Gal4* flies were crossed with the isogenic control line VDRC# 60000, generating the genotype *brp-GFP / +, repo-Gal4 / +*). Three samples with three heads each were collected from each cross. Heads were homogenized and GFP protein concentration measured from these lysates using an ELISA assay. (B) For each gene, the fold change and p-value were calculated relative to controls. On the left (red), are genes that, when knocked down in glia, reduce Brp-GFP levels and those to the right (green), are genes that when knocked down, increase Brp-GFP. (C) Hit criteria were defined as genes that when knocked down resulted in a >25% fold-change in Brp-GFP levels with a p<0.01, and genes with a >15% change and p<0.05 were also identified. (D) Gene ontology analysis on all identified genes was performed and the top 10 significantly enriched biological pathways associated with genes whose knockdown increased Brp-GFP and (E) genes whose knockdown decreased Brp-GFP were determined.

We then used our collection of RNAi lines to knockdown each of the selected genes, one at a time, specifically in glia using the *repo-Gal4* driver and evaluated Brp-GFP levels by ELISA (Fig 1B). Of the genes tested, 45 resulted in lethality when knocked out in glia and Brp-GFP levels were assessed in the remaining 926 lines (Fig 2B). In total, we found 48 genes that, when knocked down in glia, resulted in a 25% or greater change in Brp-GFP levels with a significance value cut-off of p<0.01, and an additional 135 that resulted in >15% changes with p<0.05 (Fig 2C).

To gain insight into the general categories of genes involved in glial regulation of synapse development, we performed gene ontology analysis on all of the genes identified in the screen that, when knocked down, resulted in a >15% increase (Fig 2D, S5A-B) or decrease (Fig 2E, S5C-D) in Brp-GFP levels. Both included networks of genes involved in regulation of synapses or neuronal development in mammals, though in many of these cases, glia had not been previously implicated in the action of these genetic pathways. Among those genes whose loss increased Brp-GFP levels, enriched pathways included cell adhesion and ECM regulatory molecules, immune molecules and other signaling pathways (Fig 2D, S5A-B). Those genes that, when knocked down, decreased Brp-GFP levels were enriched in several metabolic and catabolic processes (Fig 2E, S5C-D). Several screen hits were further validated by both ELISA (Fig S6A) and imaging (Fig S6B-D). Importantly, some of the categories molecules identified, such as neurotransmitter receptors, would likely only have been identified in an *in vivo* screen, where glia maintain their endogenous interactions with neurons and those neurons are engaging in normal patterns of activity. Altogether, the results of this screen demonstrate the breadth of molecular pathways glia use to regulate synapse development, and provide molecular footholds to start dissecting these mechanisms.

### Loss of glial Crq increases synapse numbers in development

Previous studies have identified immune-related molecules – complement proteins^15^, chemokines^39^ and other immune receptors^17^ – as important regulators of synapse development in mammals. In line with this previous work, the most enriched pathway among the negative regulators of synapse development was pattern recognition receptors, part of a large network of pathways related to immune function. Specifically, we identified a class of immune molecules – scavenger receptors – six of which resulted in altered synaptic protein levels during development. To begin to explore the role that scavenger receptors might play in synapse development, we chose to focus on one of these, Croquemort (Crq), which also produced highly consistent results in our secondary screening. This two-pass transmembrane receptor is orthologous to two mammalian scavenger receptors, *CD36* and *SCARB2. SCARB2* mutations are the genetic cause of a rare, progressive form of epilepsy^51^, demonstrating that this gene is somehow important for regulating circuit development, though the cause of this phenotype is unknown. Furthermore, both *SCARB2* and *CD36* variants have been associated with other diseases characterized by pathological synaptic changes, including autism^52^, Alzheimer’s disease^53,54^ and Parkinson’s disease^55,56^. These genetic associations suggest that Crq may have a conserved role in regulating synaptic circuits, and thus we chose to move forward with validating and understanding the function of glial Crq in synapse development.

Because the screen was based on measuring levels of a synaptic molecule, using an ELISA-based measurement of the synaptic protein Brp, we first wanted to validate these findings using orthogonal methods for measuring synapses, and additional genetic targeting strategies to manipulate Crq expression (Fig S7A). Two RNAi lines in the screen that targeted *crq* resulted in an increase in Brp-GFP levels. Two additional RNAi lines were available from public repositories, and we found that knockdown of *crq* using any of these four RNAi lines resulted in a significant increase in Brp-GFP levels (Fig 3A). As with knockdown of *crq* in glia, we found that *crq* mutants^57,58^ exhibited significant and substantial increases in Brp protein levels by Western blot (Fig 3B, C). These changes in Brp level could also be rescued by re-expression of *crq* in glia (Fig 3D, E), comparable to levels when *crq* was re-expressed in all Crq-expressing cells (Fig S7B, C). In addition to Brp, *crq* knockdown in glia also resulted in increases in levels of an additional presynaptic protein, Syt7^59^ (Fig 3F) and a postsynaptic protein, Dlg1^60^ (Fig 3G). These results indicate that Crq in glia coordinately regulates multiple synaptic proteins, and therefore likely synapses themselves. To further address this, synapses evaluated by imaging and an analysis of mean Brp intensity across all of the synaptic regions of the *Drosophila* brain performed (Fig S7D). In almost all regions, knockdown of crq in glia produced a significant increase in signal intensity. Thus, Crq acts to regulate synaptic protein levels broadly across the brain. To evaluate whether this was due to a change in the number of presynaptic structures, we counted the number of Brp^+^ puncta in the antennal lobes (Fig 3H). We observed an increase in puncta number in this region (Fig 3I), suggesting that an increased number of synapses is at least partially responsible for the increase in synaptic protein levels we observed.

**Figure 3:**
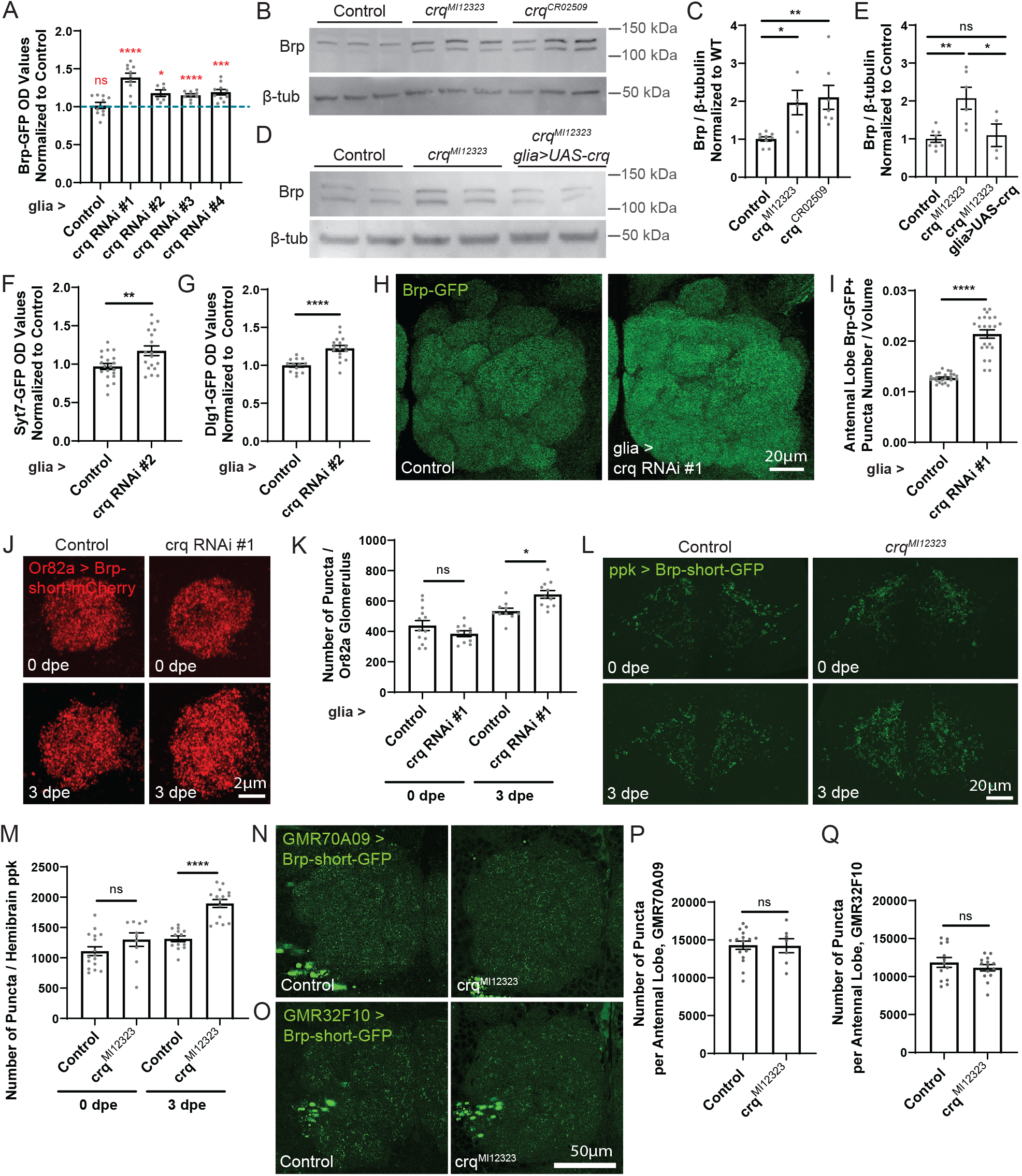
Loss of glial Crq increases synapse numbers in development. (A) Brp-GFP levels were measured from samples in which 3 heads of each of the indicated genotypes were pooled and homogenized and then analyzed by ELISA. Each of the available RNAi lines was used to knockdown crq in glia in flies with the genotype *brp-GFP / +, repo-Gal4 / + and heterozygous for UAS-crq RNAi#1* (VDRC#45884), *UAS-crq RNAi #2* (VDRC#45883), *UAS-crq RNAi#3* (BL#40831), *UAS-crq RNAi #4* (BL#42811). Results were normalized to the average value of controls (*brp-GFP, repo-Gal4* flies crossed with the isogenic control line VDRC# 60000, dotted line). Males and females at 3 dpe were used. The n for each group are: 10 (control), 10 (crq RNAi #1), 6 (crq RNAi #2), 10 (crq RNAi #3), 10 (crq RNAi #4). All groups are normally distributed. A one sample t-test was performed using against a hypothetical mean of 1 for each sample. (B) Brp-GFP protein levels were evaluated by Western blot in two crq mutant lines, *crq*^*MI12323*^ and *crq*^*CR02509*^ and *yw* controls. Five heads were pooled from male and female flies at 3 dpe for each sample. (C) The number of samples analyzed for each group (n) are: 9 (control), 4 (*crq*^*MI12323*^), 7 (*crq*^*CR02509*^). Brp levels were normalized to β-tubulin for each sample. Data from all groups are normally distributed. Samples were compared using an ordinary one-way ANOVA (**) and Sidak’s post-hoc comparisons indicated. (D) Western blots were performed on 5 pooled heads per sample from 3 dpe male and female flies for the indicated genotypes. (E) The number of samples analyzed in each group are: 8 (control – *yw*), 6 (*repo-Gal4 / +, crq*^*MI12323*^), and 4 (*repo-Gal4 / +, UAS-crq / +, crq*^*MI12323*^). Samples were compared using an ordinary one-way ANOVA (**) with Tukey’s post-hoc tests indicated. (F) GFP protein levels were assessed by ELISA from flies expressing endogenously tagged Syt7-GFP. Three heads were pooled for each sample for 3 dpe control (*syt7-GFP / +, repo-Gal4 / +*) and crq RNAi #2 (*syt7-GFP / +, repo-Gal4 / +, UAS-crq RNAi #2 / +*) flies and GFP concentration evaluated by ELISA. Males and females were run separately and the samples from each sex were normalized to same-sex controls, and then male and female data were combined. The n for each group are: 20 (control) and 18 (crq RNAi#2). Data from both groups are normally distributed. The results from an unpaired t-test are indicated. (G) GFP levels were also analyzed by ELISA for flies expressing Dlg1-GFP (control - *dlg1-GFP* / + (⍰) *or dlg1-GFP / Y* (⍰), *repo-Gal4* / + and crq RNAi#2 -*dlg1-GFP* / + (⍰) or *dlg1-GFP* / *Y* (⍰), *repo-Gal4* / *+, UAS-crq*-*RNAi#2 / +*). Three fly heads were pooled for each sample. Samples from 3 dpe male and female flies were run separately and the samples from each sex were normalized to the average value of same-sex controls. These data were then combined. The n for each group are: 13 (control) and 16 (*crq RNAi#2*). Data from both groups are normally distributed. Groups were compared using an unpaired t-test and results are indicated. (H) Endogenous Brp-GFP signal was imaged in the antennal lobe from control (*brp-GFP / +, repo-Gal4 / +*) and glia > crq RNAi#1 (*brp-GFP / +, repo-Gal4 / +, UAS-crq RNAi#1 / +*) male and female flies. Images were acquired using a 63x objective with 0.2μm z steps across the entire antennal lobe (∼25μm total depth). Representative images are z projections through the entire antennal lobe. (I) The density of Brp^+^ puncta number evaluated by counting the number of puncta in 3D through the entire antennal lobe and normalizing it to total antennal lobe volume for each brain. This was performed in male and female flies at 3 dpe from control (*brp-GFP / +, repo-Gal4 / +*) and crq RNAi#1 (*brp-GFP / +, repo-Gal4 / +, UAS-crq RNAi #1 / +*) groups. Each n represents the results from an antennal lobe from one individual fly. The n for each group are: 27 (control) and 22 (crq RNAi#1). Data from the control group are normally distributed but data from the crq RNAi#1 group is not, so they were compared using an unpaired, nonparametric Mann-Whitney test. (J) Synapses were labeled with Brp-short-mCherry in Or82a neurons, a subset of neurons which project to a single glomerulus within the fly antennal lobe (control – *Or82a-lexA / +, lexAop-brp-short-mCherry / +, repo-Gal4 / +;* crq RNAi#1 - *Or82a-lexA / +, lexAop-brp-short-mCherry / +, repo-Gal4 / +, UAS-crq RNAi#1 / +*). One glomerulus was imaged from each brain from male and female flies at the time points indicated. Images were acquired using a 63x objective with 0.25μm z steps through the entire glomerulus and z projections through this volume are shown in the representative images. (K) The number of synapses across the volume of the glomerulus were quantified. This analysis was performed as in (I) above. Each n represents a glomerulus from one individual fly. The n for each group are: 13 (control, 0 dpe), 10 (crq RNAi#1, 0 dpe), 10 (control, 3 dpe), 11 (crq RNAi#1, 3 dpe). All groups are normally distributed and compared using a two-way ANOVA (age ****, genotype – ns, interaction **). Sidak’s post-hoc tests are shown. (L) Synapses within ppk-expressing neurons were visualized using Brp-short-GFP in male and female control (*ppk-Gal4 / +, UAS-brp-short-GFP / +*) and *crq*^*MI12323*^ mutants (*ppk-Gal4 / +, UAS-brp-short-GFP / +, crq*^*MI12323*^) at the time points indicated. Images were acquired using a 40x objective with z sections 0.22μm apart, acquired across the ventral neuropil region through the depth in which ppk synapses were labeled (∼35μm). Representative images show z projections across the entire ventral neuropil. (M) The number of GFP^+^ puncta per hemibrain was quantified using the “spots” function in Imaris to identify individual puncta across the volume of the region. Each n represents a hemibrain analyzed from one individual fly. The n for each group are: 16 (control, 0 dpe), 10 (crq^MI12323^, 0 dpe), 13 (control, 3 dpe), 15 (crq^MI12323^, 3 dpe). All groups were normally distributed, except for the crq^MI12323^, 0 dpe group. Groups were compared using a two-way ANOVA (genotype ****, age ****, interaction *) and results from Sidak’s post-hoc tests are shown. (N) Synapses were labeled in inhibitory LN neurons using two separate drivers: GMR70A09-Gal4 (control - *GMR70A09-Gal4 / +, UAS-brp-short-GFP /* + and crq^MI12323^ *– GMR70A09-Gal4 / +, UAS-brp-short-GFP / +, crq*^*MI12323*^) and (O) GMR32F10-Gal4 (control – *GMR32F10-Gal4 / +, UAS-brp-short-GFP / +* and crq^MI12323^ – *GMR32F10-Gal4 / +, UAS-brp-short-GFP / +, crq*^*MI12323*^). In both cases, male and female flies were used at 3 dpe. Images were acquired across the entire antennal lobe using a 63x objective with z slices 0.2μm apart across a depth of ∼25μm. Representative images are z projections. The larger green spots in the lower left-hand corner of the images are GFP signal associated with nearby neuronal cell bodies and were excluded from the analysis. (P) The number of puncta within an antennal lobe for each brain was quantified within the GMR70A09 labelled population. Each n represents this value for an individual fly. The n for each group are: 16 (control) and 7 (crq^MI12323^). Data from both groups were normally distributed and compared using an unpaired t-test. (Q) Puncta were similarly quantified across the antennal lobe within the GMR32F10 labelled population. The n for each group are: 13 (control) and 14 (crq^MI12323^). Data were normally distributed and were compared using an unpaired t-test.

A unique advantage of using *Drosophila* to study synapse formation and elimination is that we can monitor these processes in desired subsets of neurons. Using existing models, the field has been limited in the subtypes of neurons which can be studied based on the small number of neuronal subtypes that can be successfully cultured *in vitro* without glia, and *in vivo* to subsets of excitatory neurons in primary sensory areas that can be clearly labeled and manipulated. In contrast, there are thousands of driver lines that allow cell-type specific visualization and manipulation of different neuronal subsets in the fly^61^, and these can be used to express synaptic labels specifically in those cells^62^. To evaluate the effect of manipulating Crq expression on synapses in specific populations of neurons, we expressed a synaptic label in three populations: excitatory Or82a-expressing neurons that are part of the olfactory system^63^, neuromodulatory ppk neurons, which regulate thirst behavior in the adult brain^64^ and two subsets of inhibitory lateral interneurons which are involved in higher level processing of olfactory signals. In Or82a neurons, knockdown of *crq* in glia resulted in an increase in presynaptic puncta at 3 dpe, consistent with what we observed when assaying synapses globally (Fig 3J, K). Interestingly, these differences were not present at 0 dpe (Fig 3J, K). This supports the notion that these Crq-dependent changes were acquired during the dynamic period of synaptic remodeling that we found occurs during early adulthood, rather than the time of initial circuit formation. Similar to what we observed in Or82a^+^ neurons, crq mutants did not exhibit differences in synapse number compared to controls in ppk^+^ neurons at eclosion, but a substantial increase presynaptic puncta in *crq* mutant flies was evident by 3 dpe (Fig 3L, M). In contrast, loss of Crq had no effect on synapse number within inhibitory lateral interneurons (Fig 3N-Q). This suggests that there may be some specificity to the types of synapses regulated by Crq.

*Crq* mutants and *crq* knockdown in all glia resulted in an increase in synapses. To determine the glial subtype responsible for these changes, we first assessed in which cell types *crq* is expressed using a *crq* transcriptional reporter to drive a nuclear label (Fig S8A). When co-staining with a marker for all glial nuclei (repo), we found that >90% of cells expressing *crq* were glia and this was consistent across all ages (Fig S8B). The *crq* transcriptional reporter also showed a decrease in the levels of *crq* expression over developmental time (Fig S8C).

Next, we knocked down *crq* in different subtypes of *Drosophila* glia: astrocytes, the glial subtype in direct contact with most synapses in the *Drosophila* brain; cortex glia, which are associated with neuronal cell bodies; and ensheathing glia which sit in between those two regions^65^. As expected, knockdown of *crq* just in astrocytes produced a significant increase in Brp-GFP levels, while crq knockdown in ensheathing glia did not (Fig S8D). Surprisingly, knocking down crq in cortex glia also resulted in an increase in Brp-GFP levels (Fig S8D), even though these cells have no direct contact with synapses. One possible explanation could be that, in addition to selective elimination of synapses, whole neuron elimination could contribute to the decrease in synapses observed during development, and perhaps knockdown of *crq* in cortex glia might prevent the elimination of neuron cell bodies (and their associated synapses). When we labeled neuronal nuclei (elaV) and co-stained with an antibody that recognizes cleaved Dcp1 to label dying cells, we found no instances of cleaved Dcp1^+^elaV^+^ cells (Fig S9A). There were only up to 3 total cleaved Dcp1^+^ cells per brain during the time points when synapse elimination occurs, and no difference in these numbers was observed when *crq* was knocked down in glia (Fig S9B). However, when we counted the number of neuronal cell bodies, we did find that there was a small, but consistent reduction in neuronal nuclei counts between 3 and 10-12 dpe (Fig S9C-H). To test whether Crq was required for this process, we evaluated whether this neuronal loss was attenuated when crq was knocked down in all glia (Fig S9C), just in astrocytes (Fig S9D) or just in cortex glia (Fig S9E). However, Crq was not required for developmental neuron loss to occur in any of these conditions (Fig S9F-H). This suggests that the retention of synapses observed with loss of Crq is not simply due to retention of supernumerary neurons, and also that genetically separable mechanisms regulate elimination synapses versus neuronal cell bodies.

### Crq regulates synaptic engulfment

The increase in synapses observed with loss of Crq could be due to increased synapse formation or decreased synapse elimination. Based on Crq’s known role in debris clearance in the immune system^66^, we anticipated that Crq would be involved in synapse elimination. To assess this, we first evaluated whether knockdown of *crq* affected levels of the genetically encoded engulfment markers, FYVE-GFP (Fig 4A, B) and spn-RFP (Fig 4A, C) that are expressed in astrocytes at high levels at 3 dpe. Indeed, we found that knockdown of crq in astrocytes strongly reduced these markers of engulfment. To ensure that *crq* knockdown did not simply interfere with the genetic expression of these constructs, we also used a non-genetically encoded marker of engulfment activity, LysoSensor, which labels acidic lysosomes in live brain tissue (Fig S10A). Similar to the other engulfment markers, *crq* knockdown resulted in a significant reduction in LysoSensor signal across the brain compared to controls (Fig 4D). We did not observe any gross changes in astrocyte distribution or morphology (Fig S10B, C), and therefore the changes we see in astrocyte engulfment markers are not likely due to diminished astrocyte number or infiltration. Together, these results argue that Crq is required for the phagocytic activity of glia during early adulthood.

**Figure 4:**
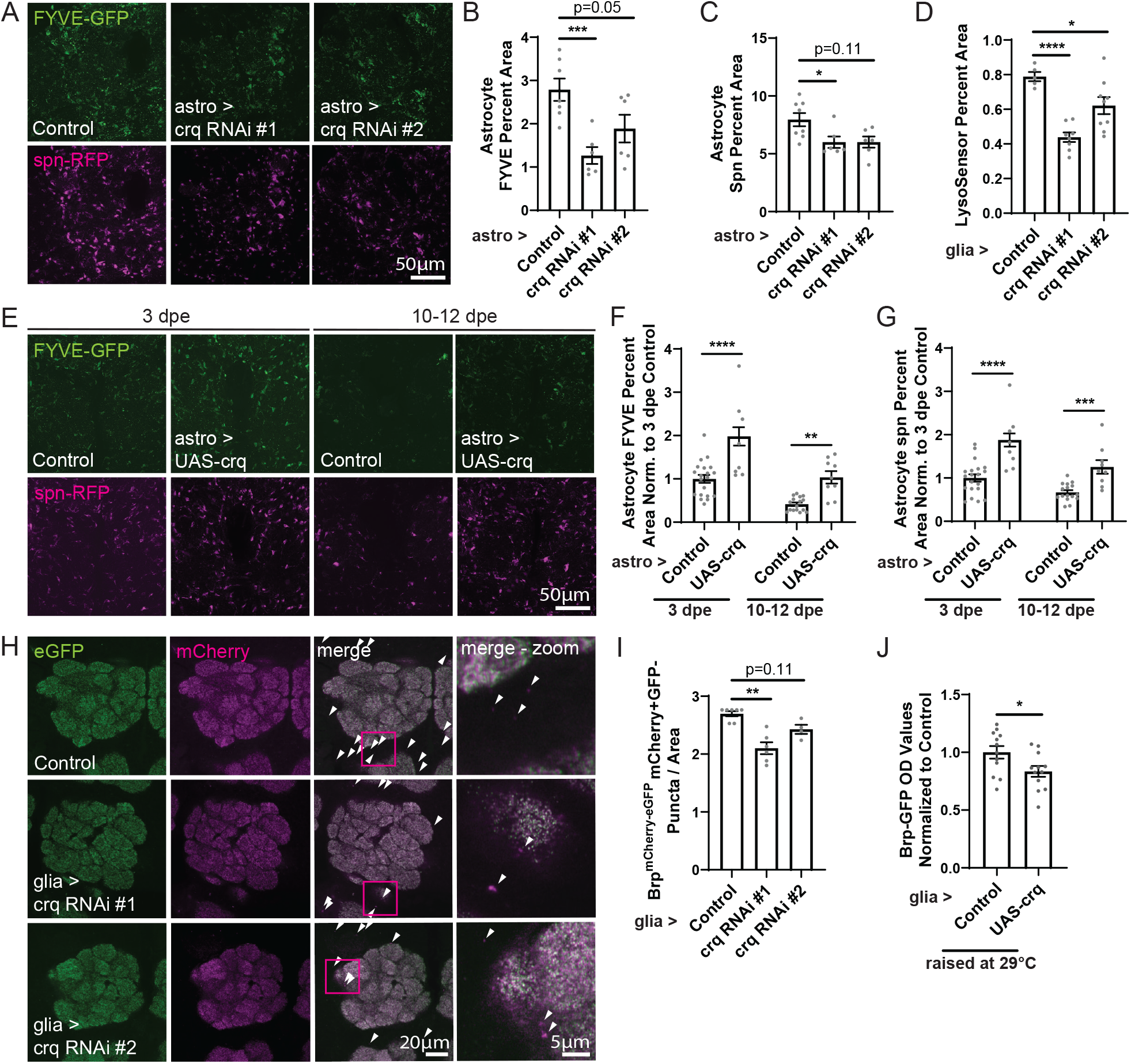
Crq regulates glial synaptic engulfment. (A) FYVE-GFP and spn-RFP engulfment indicators were expressed astrocytes (control – *GMR25H07-Gal4 / +, UAS-FYVE-GFP/+, UAS-spn-RFP/+* and *GMR25H07-Gal4 / +, UAS-FYVE-GFP/+, UAS-spn-RFP/+, 10xUAS-FLP;* astro>crq RNAi#1 - *GMR25H07-Gal4 / +, UAS-FYVE-GFP/+, UAS-spn-RFP/+, UAS-crq RNAi#1 / +;* astro>crq RNAi#2 - *GMR25H07-Gal4 / +, UAS-FYVE-GFP/+, UAS-spn-RFP/+, UAS-crq RNAi#2 / +*). Male and female flies were collected at 3 dpe. Images were acquired with a 40x objective and the central brain was imaged starting at the anterior-most portion and z slices acquired every 5μm through a total depth of 45μm. Representative images show z projections through this depth in a portion of the central brain. (B) Z projections were made and the percent FYVE+ area was quantified across the central brain. Each n represents this percentage for one brain. The n for each group are: 7 (control), 6 (crq RNAi#1) and 6 (crq RNAi#2). All data are normally distributed. Groups were compared using an ordinary one-way ANOVA (**) with Sidak’s post-hoc comparisons indicated. (C) The percent spn+ area was quantified as described in (B). The n for each group are: 8 (control), 6 (crq RNAi#1) and 6 (crq RNAi#2). control and crq RNAi#2 groups were normally distributed, but the crq RNAi#1 group was not. A Kruskal-Wallis test was used to compare data across these groups (*) and results of Dunn’s post-hoc tests are shown. (D) LysoSensor was applied to acutely dissected male and female fly brains at 3 dpe. Images were acquired using a 40x objective and 3 z planes were taken 5μm apart. For each slice, the neuropil regions within the central brain were outlined and the percent LysoSensor+ area within those regions quantified. This value was averaged across the 3 z planes. Each n represents one individual brain. The n for each group are: 5 (control *-repo-Gal4 / +*), 8 (crq RNAi#1 *– repo-Gal4 / +, UAS-crq RNAi#1 / +*), and 9 (crq RNAi#2 *– repo-Gal4 / +, UAS-crq RNAi#2 / +*). Data from all groups are normally distributed. Data were analyzed using an ordinary one-way ANOVA (****) and Dunnett’s post-hoc test results are shown. (E) FYVE-GFP and spn-RFP were expressed in astrocytes and the signal assessed in controls compared to flies in which an astrocyte driver is used to express UAS-crq in male and female flies at the time points indicated (control *– GMR25H07-Gal4 / +, UAS-FYVE-GFP/+, UAS-spn-RFP/+*) and UAS-crq (*GMR25H07-Gal4 / +, UAS-FYVE-GFP/+, UAS-spn-RFP/+, UAS-crq / +*). Images were acquired using a 40x objective and z sections were acquired every 5μm apart from the anterior-most portion of the central brain to a depth of 45μm. (F) The percent FYVE+ area was assessed across the central brain from 3 z slices, starting with the first section where the antennal lobes were both clearly distinguishable and every 10μm from that point. The percent areas were averaged across these 3 images. These values were then normalized to the average value for the 3 dpe control group. Each n represents one brain. The n for each group are: 20 (control, 3 dpe), 13 (UAS-crq, 3 dpe), 16 (control, 10-12 dpe), 9 (UAS-crq, 10-12 dpe). All groups are normally distributed, except for the 10-12 dpe control group. Data were compared using a two-way ANOVA (age ****, genotype ****, interaction – ns) and Sidak’s post-hoc tests are shown. (G) Spn+ area was quantified as described in (F). The n for each group are: 20 (control, 3 dpe), 13 (UAS-crq, 3 dpe), 16 (control, 10-12 dpe), 9 (UAS-crq, 10-12 dpe). Data from all groups are normally distributed. Data were compared using a two-way ANOVA (age ****, genotype ****, interaction – ns) and Sidak’s post-hoc tests are indicated. (H) Brp was endogenously labeled with a pH-sensitive eGFP and a pH-insensitive mCherry in male and female flies at 3 dpe in control (*brp*^*mCherry-eGFP*^ */ +, repo-Gal4 / +*), glia> crq RNAi#1 (*brp*^*mCherry-eGFP*^ */ +, repo-Gal4 / +, UAS-crq RNAi#1 / +*) and glia> crq RNAi#2 (*brp*^*mCherry-eGFP*^ */ +, repo-Gal4 / +, UAS-crq RNAi#2 / +*) groups. Brains were imaged with a 40x objective and z planes were acquired starting at the anterior-most portion of the brain and every 5μm for 8-10 z planes. The signal was overlayed (merge) and mCherry+eGFP-puncta were identified (indicated by white arrows, shown magnified in merge-zoom insets). (I) The number of mCherry+eGFP-puncta were quantified across the central brain in each z section and then normalized to the total area quantified. These values were then averaged across each brain. Each n represents one brain. The n for each group are: 7 (control), 6 (crq RNAi#1) and 4 (crq RNAi#2). Data in the control group are not normally distributed, and data for the crq RNAi#1 and crq RNAi #2 groups are. Data were compared using a Kruskal-Wallis test (***) and Dunn’s post-hoc tests are indicated. (J) Male and female flies (control *– brp-GFP / +, repo-Gal4 / +* and UAS-crq – *brp-GFP / +, repo-Gal4 / +, UAS-crq /* +) were raised at 29°C starting at eclosion, and were collected at 3 dpe. Two fly heads were pooled per sample and GFP concentrations from each sample measured by ELISA. The n for each sample are: 12 (control) and 12 (UAS-crq). Data from both groups are normally distributed. Data were compared using an unpaired t-test.

Using a *crq* transcriptional reporter, we had seen that *crq* expression begins to decrease after 3 dpe (Fig S3D). We wondered whether this reduction in *crq* expression might actually cause the decrease in glial engulfment observed between 3 dpe and 10-12 dpe. To test this, we generated a fly in which *crq* is expressed under UAS control so that we could overexpress *crq* using an astrocytic Gal4 driver, not subject to its normal patterns of transcriptional regulation across development. We found that *crq* overexpression in astrocytes resulted in an increase in both FYVE-GFP (Fig 4E, F) and spn-RFP (Fig 4E, G) engulfment indicators at 3 dpe. This increase was maintained out to 10-12 dpe, suggesting that *crq* expression is sufficient to drive continued glial phagocytic activity, even past the developmental time to which this process is normally restricted.

To further explore whether Crq regulated glial engulfment of synapses, we used the Brp^mCherry-eGFP^ flies we generated and evaluated the number of mCherry^+^eGFP^-^ puncta, indicative of synapses which have undergone engulfment (Fig 4H, I). Indeed, we found the knockdown of crq in glia resulted in significant reductions in mCherry^+^eGFP-signal (Fig 4I), in addition to decreases in the ratio of mCherry to eGFP positive signal (Fig S10D-F), arguing that Crq is required for normal levels of developmental synaptic engulfment.

The reduction in markers of engulfment activity when *crq* was knocked down correlates with increases in levels of Brp-GFP synaptic protein levels. We wondered whether the increases in engulfment markers observed when *crq* was overexpressed would conversely result in a reduction in Brp-GFP levels. To test this, we measured Brp-GFP levels by ELISA, and found that strong overexpression of crq did result in a reduction in synaptic protein (Fig 4J). Taken together, this demonstrates that loss of Crq is sufficient to reduce glial engulfment, and overexpression of Crq is sufficient to enhance glial engulfment activity.

### Crq is required for developmental and age-related synapse elimination and proper circuit function

If Crq regulates synaptic engulfment, we would expect synapse elimination to be attenuated in flies in which *crq* is knocked down in glia. When we established the time course of synapse development in the fly, we had identified that synaptic protein levels decrease between 3 and 10-12 dpe (Fig 1C). When we knocked down *crq* in glia, this reduction in synapses during development no longer occurred (Fig 5A), indicating that Crq is required for developmental synapse elimination.

**Figure 5:**
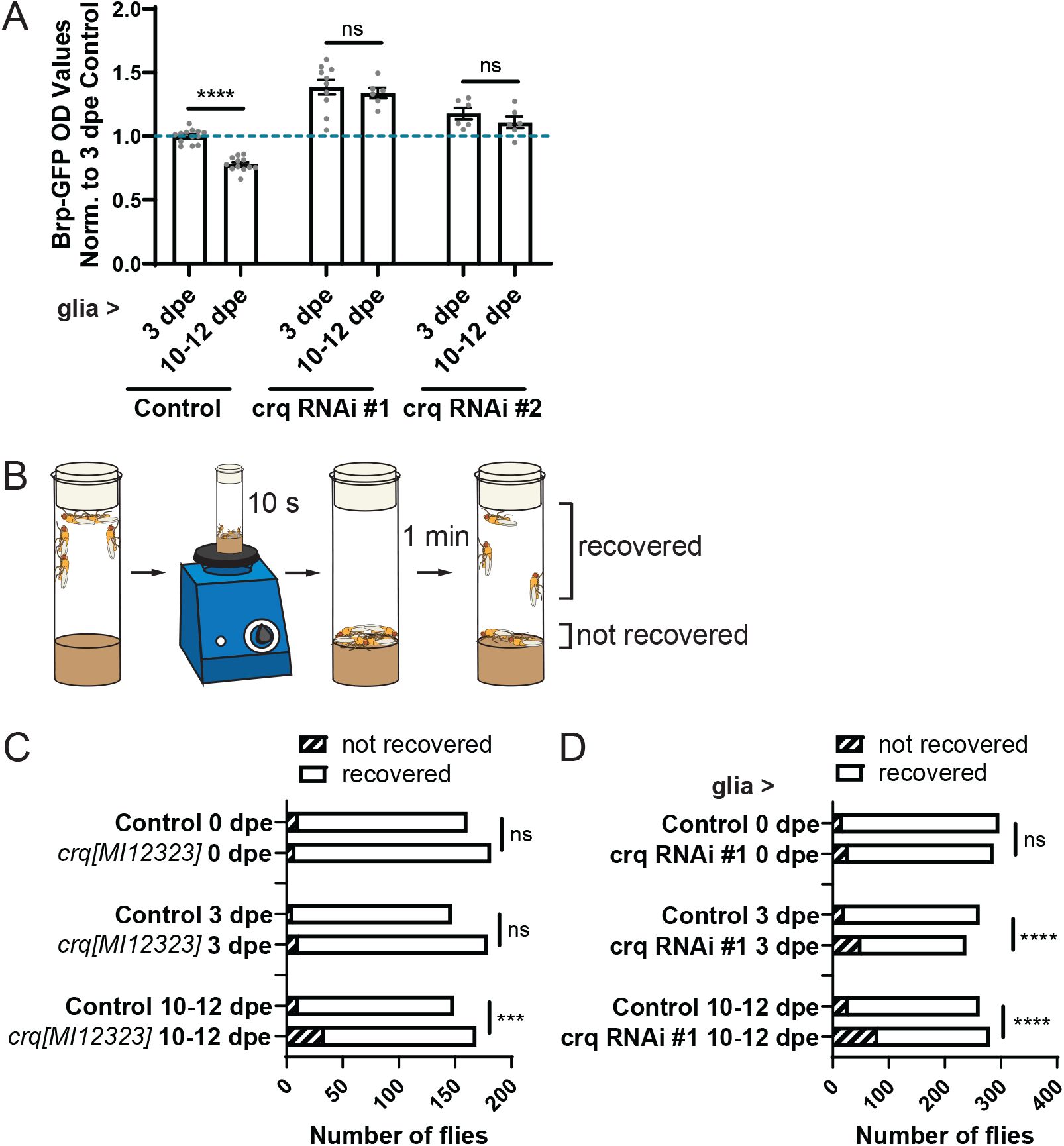
Crq is required for developmental synapse elimination and proper circuit function. (A) Brp-GFP levels were quantified in fly head lysates by ELISA across development. Two fly heads were pooled for each sample for control (*brp-GFP / +, repo-Gal4 / +*), crq RNAi#1 (brp-GFP / *+, repo-Gal4 / +, UAS-crq RNAi#1 / +*) and crq RNAi#2 (*brp-GFP / +, repo-Gal4 / +, UAS-crq RNAi#2 / +*) groups. Male and female flies were run separately and normalized to their own 3 dpe control average. The n for each group are: 13 (control, 3 dpe), 12 (control, 10-12 dpe), 10 (crq RNAi#1, 3 dpe), 6 (crq RNAi#1, 10-12 dpe), 6 (crq RNAi#2, 3 dpe), 6 (crq RNAi#2, 10-12 dpe). Data from all groups are normally distributed. These groups were compared using a two-way ANOVA (age ***, genotype ****, interaction *). Results from Sidak’s post-hoc tests are shown. (B) A seizure-susceptibility assay was used in which flies are vortexed for 10s and recovery scored by evaluating the number of flies that resumed normal climbing behavior after 1 min. (C) Recovery from vortexing was scored in male and female controls (yw) and crq^MI12323^ mutants. Chi-squared tests were used to evaluate differences between these two genotypes at the developmental stages indicated. (D) Recovery from vortexing was scored in male and female control (*repo-Gal4 / +*) flies and flies in which *crq* was knocked down in glia (*repo-Gal4 / +, UAS-crq RNAi #1 / +*) at the time points indicated.

We reasoned that retention of supernumerary functional synapses within fly circuits should affect circuit function. Given our findings that excitatory, but not inhibitory synapse number was changed with loss of glial *crq*, and that human mutations in *crq’s* ortholog, SCARB2, are the cause of a progressive form of epilepsy^51^, we evaluated whether *crq* mutants might also exhibit enhanced circuit activity. To test this, we used an assay to assess seizure susceptibility (Fig 5B). The strong sensory stimulus of vortexing flies can result in seizures in flies with heightened circuit excitability, resulting in transient immobility and impaired climbing behavior^67^. In *crq* mutant flies, there was no difference in recovery at 0 dpe, but by 10-12 dpe there was a significant increase in the proportion of mutant flies that failed to recover within 1 minute relative to controls (Fig 5C). Knocking down *crq* in glia produced similar results, with no difference in seizure susceptibility at 0 dpe, a significant increase by 3 dpe, and a further enhancement of this behavior by 10-12 dpe (Fig 5D). The time course of these behavioral changes aligns well with when increased synaptic protein levels were observed in these flies, and with the progressive epilepsy phenotype observed in humans with *SCARB2* mutations. Baseline climbing behavior was impaired in *crq* mutant flies by 10-12 dpe (Fig S11A), but not when *crq* was knocked down in glia (Fig S11B) – though a climbing deficit could be induced by enhancing the strength of glial *crq* knockdown (Fig S11C) – suggesting that general locomotor deficits are unlikely to account for the measures used to evaluate seizure recovery. Together, these data support the notion that the synaptic increases evident with loss of *crq* result in changes in brain circuitry and behavior. Like changes in synapses, these phenotypes were not present at eclosion, indicating that these effects are driven by changes in synaptic circuitry that occur in the adult fly.

Finally, increased circuit excitability has been shown to impair survival in flies, so we sought to evaluate whether loss of *crq* would impact the fly lifespan^68^. *Crq* mutant flies have been shown to exhibit early lethality due to impaired immune function^69^, which we recapitulated here (Fig S11D). However, we also found that knockdown of *crq* in glia also produced survival deficits, albeit much milder than those in *crq* mutants (Fig S11E). If these survival deficits were linked with the increased circuit excitability we observed, we reasoned that we should be able to aggravate this survival deficit in flies which already exhibit increased neuronal activity, such as in *Sh*^5^ (*shaker*) mutants^70^. Indeed, we see that, while *Sh*^5^ flies exhibit no survival deficit on their own, this mutant significantly worsens survival defects in flies in which *crq* is knocked down in glia (Fig S11E). Together, these data suggest that the synaptic changes observed in flies lacking *crq* result in progressive circuit dysfunction over the lifetime of the organism.

These long-term behavioral results lead us to test whether glial *crq* might continue to play roles outside of the context of development. In our assessment of the kinetics of synaptic changes across the *Drosophila* lifespan, we had also observed a decrease in Brp-GFP signal evident between 10-12 and 30-35 dpe, which suggests *Drosophila* experience a period of age-related synaptic loss (Fig 1C). Since knockdown of crq prevented the decrease in synapses we observed during development, we asked whether it might also attenuate this synaptic decrease associated with age. Remarkably, *crq* knockdown in glia also completely prevented this age-related decrease in Brp-GFP protein levels (Fig 6A). This attenuation of age-related synaptic loss was recapitulated when crq was knocked down in all glia (Fig S12A), just in astrocytes (Fig S12B) and just in cortex glia (Fig S12C). This complements existing work in mammals demonstrating a necessary role for glia in mediating age-related synaptic loss.^71^ We observed strong increases in markers of endosomal (Fig 6B, C) and phagolysosmal (Fig 6B, D) activity with age, and an increase in measures of synaptic engulfment (Fig 6E-G).

**Figure 6:**
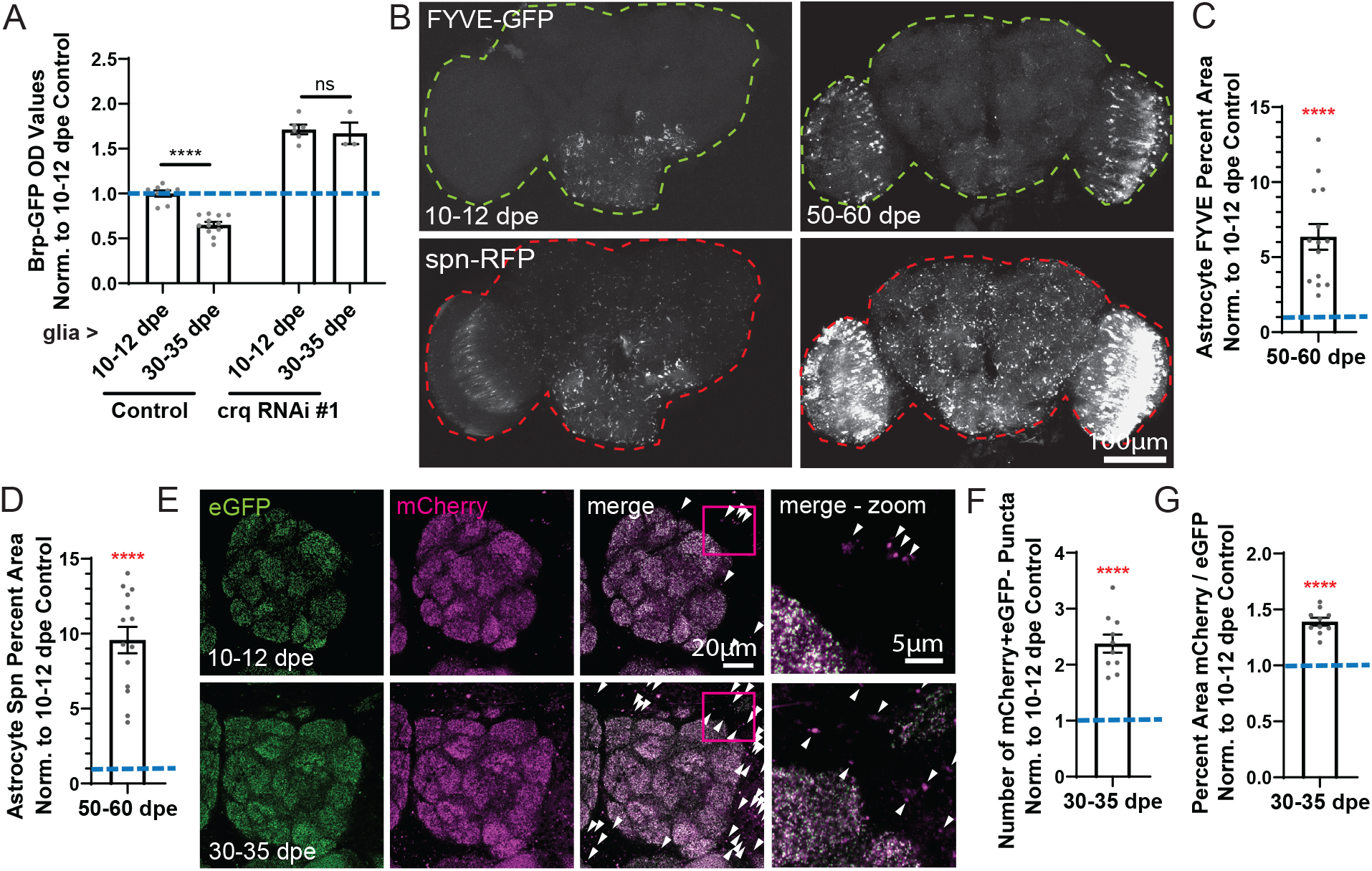
Crq is required for age-related synapse elimination. (A) Brp-GFP levels were measured by ELISA to assess changes in synaptic protein levels with age in controls (*brp-GFP / +, repo-Gal4 / +*) and when crq was knocked down in glia (crq RNAi#1 *-brp-GFP / +, repo-Gal4 / +, UAS-crq RNAi#1 / +*) in male and female flies. Two fly heads were pooled for each sample. The OD values for each sample were normalized to the average of the 10-12 dpe control group. The number of samples for each group are: 8 (control, 10-12 dpe), 12 (control, 30-35 dpe), 6 (crq RNAi#1, 10-12 dpe), 3 (crq RNAi#1, 30-35 dpe). Data from all groups are normally distributed except for the 30-35 dpe crq RNAi#1 group. Groups were compared using a two-way ANOVA (age ***, genotype ****, interaction **). Sidak’s post-hoc tests are shown. (B) To measure changes in markers of endosomal and phagosomal activity within astrocytes in aging flies, FYVE-GFP and spn-RFP engulfment indicators were expressed astrocytes (*alrm-Gal4/+, UAS-FYVE-GFP/+, UAS-spn-RFP / +*) with males and females combined. Images were acquired with a 10x objective, with 12 z sections, beginning at the anterior-most portion of the brain and taken 5μm apart. Images were then z projected through the entire set of z planes acquired, thresholded and (C) the percent FYVE+ area measured across the entire brain area (outlined with a green dotted line) or (D) the percent spn-RFP+ area (outlined with a red dotted line) were evaluated. These values were normalized to the average value of the 10-12 dpe group which are shown in Figure 1 (blue dotted line). The n for 50-60 dpe group is 14 and these data are normally distributed. A one sample t-test was performed against the hypothetical value of 1 and results are shown. (E) eGFP and mCherry signal were visualized across the entire central brain area of male and female *brp*^*mCherry-eGFP*^ / + flies. Brains were imaged with a 40x objective and z planes were acquired starting at the anterior-most portion of the brain and taken every 5μm for 8-10 z planes. (F) The number of mCherry^+^eGFP^-^ puncta were quantified across the central brain in each z section and the average number of mCherry+eGFP-puncta / section was calculated. This value was then normalized to the average for the 10-12 dpe group shown in Figure 1 (blue dotted line). Each n represents this value from one individual brain. The n for the 30-35 dpe samples is 10 and these data are normally distributed. A one sample t-test was performed against the hypothetical value of 1 and results are shown. (G) The first 5 z planes of the images in E were analyzed, starting at the most anterior plane where the antennal lobes were both fully visible, and spaced 5μm apart. The percent area which were mCherry+ and eGFP+ were recorded across the central brain, and averaged across these 5 z planes, and this average value was recorded. Then these values were normalized to the average value from the 10-12 dpe group shown in Figure 1/S1 (blue dotted line), and each n represents that value from one brain. The n for the 30-35 dpe samples is 10 and data are normally distributed. A one sample t-test was performed against the hypothetical value of 1 and results are shown.

To test whether loss of *crq* might prevent age-related reductions in synapses through preventing age-related neuronal loss, we first evaluated whether neuronal loss occurred with age. We did detect some cleaved DCP1+ dying cells with age (Fig S12D, E) and unlike in development, we did detect some cleaved DCP1+ neurons (Fig S12 D, F). Indeed, when we counted neuron number, we did find that there was a significant reduction with aging (Fig SS12G, H). However, glial knockdown of *crq* did not prevent this neuron loss with age; in fact, there were very slight, but significant decreases in neuron number with age when *crq* was knocked down in glia (Fig S12I). Thus Crq is required for synaptic loss with age, but not age-related loss of neuronal cell bodies.

It is possible that Crq could be required for glia to engage in engulfment regardless of substrate or context, or rather it may specifically mediate selective synapse elimination. To test this, we assessed whether glia in which *crq* was knocked down could successfully engage in elimination of synaptic debris during metamorphosis and clearance of neuronal debris after injury. During metamorphosis, glia clear virtually all synapses from the larval nervous system in preparation to construct the nervous system of the adult fly^72^ (Fig S13A). We did not find any changes in clearance of synaptic debris in this context with glial *crq* knockdown (Fig S13B). To test whether Crq was required for clearance of neuronal debris after injury^73^, we unilaterally ablated a maxillary palp, injuring sensory neurons that project into the antennal lobes within the brain, and simultaneously labeled a subset of those neurons to visualize their projections (Fig S13C). We found that there was no difference in the extent of clearance of these projections when *crq* was knocked down in glia (Fig S13D). Together, these results demonstrate that Crq is not required for glia to engage in engulfment in all contexts, but rather is specifically required for the selective elimination of synapses that occurs in circuit remodeling during synapse development and in aging.

We do know that both clearance of synaptic debris during metamorphosis and after injury are dependent on the previously defined glial engulfment receptor Drpr^73^. We had seen that, like *crq*, knockdown of *drpr* resulted in increased synaptic protein levels in development. However, interestingly, *drpr* was not required for age-related synaptic loss to occur (Fig S13E), suggesting that Crq is unique compared to other known astrocytic phagocytic receptors in its ability to regulate age-related reductions in synaptic protein levels. These two receptors also showed distinct patterns of regulation in their expression in different contexts, with *drpr* but not *crq* increased in response to injury (Fig S13F), but both increasing with age (Fig S13G). Therefore, these receptors seem to represent parallel mechanisms to regulate synapse elimination across different contexts.

## Discussion

Synapse formation and elimination contribute to neural circuit development, and require coordination between neurons and glia. Work over the last two decades has identified a number of glial molecules that regulate synapse formation or elimination, but an emerging theme appears to be that glial regulation of synapse numbers is dynamic and mechanistically diverse. A major aim of this study was to establish *Drosophila* as a system to study conserved genes and pathways that govern broad changes in synapse formation and elimination across the brain, to complement existing *in vitro* and mouse models. We found that *Drosophila* exhibit similar synaptic dynamics over their lifespan as mammals, with periods of developmental synapse formation and elimination, as well as age-related synaptic loss. To explore the molecular basis of glial regulation of synapse numbers, we combined a rapid, ELISA-based screening method with glial-specific gene knockdown by RNAi to identify 48 genes that are conserved in glial expression from flies to mammals, and are required in vivo for normal synapse numbers. Finally, we identify the scavenger receptor Crq is required for both developmental synapse elimination and age-related synapse loss, implying that synapse loss with age is actively driven by glial cells.

While our observations ultimately require further confirmation by examining synaptic contacts by other means, and in uniquely identifiable cells over time, our results support the notion that at late pupal development and into early adulthood, synapses are dynamically changing across the *Drosophila* brain. Total synapse number peaks by 3 days after eclosion, and then is reduced over the next week. Given that loss of Crq in glia blocks this decrease in synapse numbers early in adulthood, it appears that glial cells are responsible for promoting early loss of synapses in the adult brain. Consistent with our observations, dynamic changes in synapses have been observed during this time window in response to changes in sensory input^74-76^, and these have been shown to be regulated by the glial phagocytic receptor Draper^77^.

One reason to establish *Drosophila* as a model to study glial regulation of synapse development was to be able to perform large-scale genetic screens *in vivo* to reveal the diversity of pathways required. Our ELISA-based screen identified a list of 48 genes that, when knocked down in glia, changed synaptic protein levels, reflecting the diversity of ways in which glia can regulate synapse development. Many of the genetic pathways enriched in the screen had not been previously implicated in glial regulation of synapse development. Metabolic and catabolic pathways, transporters, neurotransmitter receptors and sphingolipid regulators all represent new categories of mechanisms that influence synaptic regulation by glia. Others, such as immune-related pathways, have been previously identified as important regulators of synapse development in mammals^78^, and our screen results confirm that they represent a major category of genes involved in this process. However, it also highlights that we have not yet identified all types of immune molecules that glia use to regulate synapse development. For example, pattern recognition receptors and scavenger receptors were novel classes of molecules identified as synaptic regulators in this screen. Multiple members of the Toll receptor family were identified, as well as secreted antimicrobial peptides and molecules involved in innate immune memory. Therefore, even in the relatively well studied area of immune-related molecules, our data suggest we still have much more to discover about the diversity of ways immune molecules can be repurposed in the context of glial regulation of synapse development.

The overarching goal of this screen was to identify starting points to build our understanding of the rules that neurons and glia use to decide when and which synapses should be formed and eliminated across the lifespan. Our discovery of Crq as a glial regulator of synapse elimination reveals new insights into this process. First, we found that the timing of *crq* expression tightly correlated with that of synapse elimination. Therefore, the downregulation of *crq* expression may be a way in which glia limit when during development synapse elimination is allowed to occur at high levels. Indeed, extending expression of *crq* later in development was sufficient to promote continued expression of phagocytic markers in glia and reduced synaptic protein levels. Understanding how Crq is regulated and signals may provide further insight into how glia decide when to promote synapse elimination and tune the relative balance of synapse formation and elimination across development. Because Crq seems to act as such a potent and sustained regulator of synapse elimination, identifying its ligand would also open up the opportunity to understand how the right synapses are selected for elimination.

Crq has also provided a handle to start understanding how different glial subtypes may work together to regulate synapse development. While astrocytes are the only glial cell type within the synaptic neuropil in *Drosophila, crq* knockdown just in astrocytes or just in cortex glia was sufficient to alter synaptic protein levels, suggesting that cortex glia may coordinate with astrocytes to facilitate synapse elimination in some way. There is precedence for the phenomenon of coordination among glial cell types in regulating synapse development in mammals^79^, and future work in *Drosophila* could allow us to dissect the transcellular signaling pathway involved.

Our characterization of Crq further reveals that glia employ different receptors to mediate clearance of synapses and other neuronal debris in a context-dependent manner. While knockdown of both *crq* and *drpr* (othologous to *MEGF10*) increased levels of synaptic proteins at 3 dpe, only Crq was required for glial-mediated synaptic loss in aging, and only Drpr was required for glial engulfment of debris during metamorphosis^72^ and after injury^73^. Why would glia use distinct pathways to mediate engulfment in these different contexts? It is known that Drpr activation promotes initiation of large-scale transcriptional changes that amplify their engulfment activity^80^. Perhaps this is beneficial in situations like metamorphosis, when virtually every synapse, along with axonal and dendritic debris, needs to be rapidly cleared from the nervous system. However, this amplification of engulfment activity may be unnecessary or undesirable when fine-tuning synaptic connectivity, thus requiring a different downstream signaling mechanism. In addition to Drpr and Crq, knockdown of other immune receptors, including other members of the scavenger receptor family, also altered Brp-GFP levels in our screen. This indicates that different receptors, even from the same class, are unable to compensate for loss of the others. It is possible that they recognize different ligands, act on different subsets of synapses, or act in a convergent pathway that requires all of them to be present. Having multiple developmental checkpoints in place may give glia more power to precisely tune their engulfment, ensuring restriction of synapse elimination to appropriate times and places.

Our work supports the emerging connection between mechanisms used to mediate synaptic elimination in development and pathological synaptic loss. Here, we found that glial Crq was required for synaptic loss in aging. In mammals, there is also precedence for reuse of complement proteins in synapse elimination in development and disease^15,71,81,82^. This begs the question of whether Crq might also mediate synapse elimination in other contexts of pathology, in response to injury, neurodegeneration or immune activation. The orthologs of *crq, SCARB2* and *CD36*, have been implicated in a number of neurological diseases, suggesting that this may indeed be the case. *SCARB2* mutations are the genetic cause of a rare, progressive form of epilepsy^51^, though how these mutations cause disease is unknown. Crq mutants and *crq* knockdown in glia both confer progressive increases in seizure susceptibility over the course of development, similar to the pattern of clinical progression observed in *SCARB2* mutant carriers. Perhaps the excess retention of synapses due to impaired synaptic engulfment by glia underlies this clinical phenotype. In addition, variants in *SCARB2* and *CD36* are associated with autism^52^, Alzheimer’s disease^53,54^ and Parkinson’s disease^55,56^. Possibly the same mechanisms that these receptors use to regulate appropriate elimination of synapses in development may also contribute to synaptic pathologies that underlie these diseases. Multiple CD36 inhibitors have been approved for therapeutic use in humans, perhaps future work in mouse models of these associated diseases could evaluate whether pharmacological inhibition of these receptors could be beneficial in limiting pathological synaptic loss.

Together, our efforts using a high-throughput ELISA-based *in vivo* forward genetic screening approach in *Drosophila* has increased our knowledge of the breadth of genes glia employ to regulate synapse development, and allowed us to study Crq as a starting point to begin decoding how glia decide when and which synapses to remove. In combination with existing *in vitro* and mouse models, we now have a powerful arsenal of complementary systems to answer the fundamental question of how glia act to establish appropriate circuits in development, and address the pressing need to understand how these processes contribute to pathological synaptic changes that lead to circuit dysfunction in disease.

## Acknowledgements

We thank the entire Freeman lab for their advice and support that has shaped this work. Special thanks to Leire Abalde-Atristain, Adel Avetisyan and Ernesto Manzo for their exceptional time and effort in providing their feedback on this manuscript. We also thank Cody Call for his generous help in developing the image analysis workflow to quantify Brp puncta number. This study was supported by NIH grants F32NS117647 and K99NS133298 to TRJ, K99NS126642 to YK, and R37-NS053538 and RO1-NS112215 to MRF. We thank the Bloomington (NIH P40OD018537) and Vienna Drosophila Stock Centers for the fly stocks used in this work.

## Author contributions

Conceptualization, TRJ, YK and MRF; Methodology, TRJ; Investigation, TRJ, VOM, MKBM, VKP, YK, JC and AS; Writing – Original Draft, TRJ; Writing – Review & Editing, TRJ, YK and MRF; Supervision, TRJ and MRF; Funding Acquisition, TRJ, YK and MRF. All authors read and approved this manuscript.

## STAR Methods

### Fly stocks and maintenance

*Drosophila* were housed at 25°C in a 12 hr light-dark cycle. Except where noted below, flies were maintained on cornmeal-molasses media. For circadian experiments, no additional protocols were in place to ensure strict circadian entrainment. Larvae were collected at wandering stage 3 (wL3), pupae for experiments evaluating synaptic elimination in metamorphosis were collected at head eversion, a developmental event that occurs ∼12 hr into metamorphosis. “Late pupae” were collected at 72-96hr APF. Adults were collected at time points across the lifespan: flies denoted as “0 dpe” were collected within 24 hr of eclosion, “3 dpe” were collected within 48-72 hr of eclosion, and those the other age ranges are as stated. Existing fly lines used in this study included: *pUAST-mCD8::GFP*^*83*^, *repo-Gal4*^*84*^, *alrm-Gal4*^*85*^, *GMR25H07-Gal4*^*61*^, *per[01]*^*86*^, *UAS-pWiz-drpr RNAi7b*^*73*^, *Brp-GFP (brp*^*MI02987-GFSTF*^)^*58*^ (BL#59292), *UAS-GFP-2xFYVE* (BL#42712), *UAS-Spinster(Spn)-mRFP* (BL#42716), *UAS-tsp RNAi* (BL#34661), *UAS-sparc RNAi* (BL#40885, HMS02133), *UAS-crq RNAi#1* (VDRC#45884), *UAS-crq RNAi #2* (VDRC#45883), *UAS-crq RNAi#3* (BL#40831), *UAS-crq RNAi #4* (BL#42811), *crq[MI12323]*^*58*^ (BL#57926), *crq[CR02509]*^*87*^ (BL#93353), *Syt7-GFP* (BL#94776), *dlg1-GFP*^*58*^ (BL#59417), *Or82a-lexA, lexAop-brp*.*short*.*mCherry* (gift from Mary Logan), *UAS-brp-short-GFP*^*62*^, *GMR70A09-Gal4*^*88*^, *GMR32F10-Gal4*^*88*^, *GMR56F03-Gal4*^*89*^ (BL#39157), *Ctxglia-split-Gal4 (wrapper-Gal4DBD, Nrv2-VP16AD)*^*90*^, *Sh[5]* (BL#111), *Or85e-mCD8::GFP*^*91*^, *crq-Gal4* (BL#25041), *UAS-H2B:mCherry*.*HA* (gift from Josh Dubnau).

In this study, we generated two new fly lines: *UAS-crq* and *Brp*^*mCherry-eGFP*.^ For the *Brp*^*mCherry-eGFP*^ fly, *mCherry* was amplified from Addgene plasmid #32002 and the *eGFP* (actually *eGFP-FIAsH-StrepII-TEV-3xFlag*) amplified from DGRC plasmid #1298. These were assembled using HiFi assembly and put into the DGRC #1297 backbone. The sequence was verified using Plasmidsaurus and injected BDSC stock BL#37403 using the MiMIC injection service from BestGene. The *UAS-crq* was made by amplifying 1.4kb of crq from cDNA RE02070 and Gibson assembly used to insert this DNA into the NotI site of the *pattB-5xUAS* plasmid. This was injected by BestGene into stock BL#24749 to insert it at the ZH-86Fb attP site.

### TRAP Seq

This dataset is publically available through GEO (accession #GSE242729) and the methods for obtaining these results have been detailed previously^31^. Briefly, an eGFP-tagged L10a ribosomal subunit was expressed in astrocytes using *alrm-Gal4, elaV-Gal80*. Pupae were collected at 48-72hr after puparium formation (APF), and adults collected at 1-3 dpe, 10-12 dpe and 50-60 dpe. Two hundred flies were pooled for each sample, and 3-5 samples were evaluated at each time point. Ribsomes were precipitated with beads, and associated RNA isolated. cDNAs were prepared and the sequencing library prepared using ribosomal reduction. Sequencing was performed on a NovaSeq flow cell with 50 million templates per sample. Fastq files were assembled using bcl2fastq. Differential gene expression was evaluated using DEseq2. Heatmaps were assembled using the heatmaply package in R.

### ELISA

ELISAs were performed to detect GFP concentrations from flies expressing *Brp-GFP, Syt7-GFP, dlg1-GFP* and *GMR25H07-Gal4, UAS-mCD8::GFP* in which astrocyte membranes were labeled with GFP. 96-well plates (Immunolon 2HB, Thermo #3455) were coated with 100μl 1:5000 dilution of chicken anti-GFP antibody (Abcam #ab13970) in carbonate buffer (30mM sodium carbonate, 70 mM sodium bicarbonate, pH 9.6), sealed and left to shake overnight at 4°C. Plates were washed 4 times with 0.1% PBST and 350μl of blocking buffer (2% milk in 0.1% PBST) added, plates sealed and left to shake overnight at 4°C. Samples were prepared from flies which had been frozen and kept at -80°C until use. Heads were removed by vortexing frozen flies and heads were placed into a 1.5mL Eppendorf tube. Samples were homogenized using a hand-held homogenizer in 120μl 0.1% PBST for 5 s and placed on ice. After all samples were prepared, samples were centrifuged at 13,000 rpm for 2 min at 4°C. Plates were washed 4 times with 0.1% PBST and 100μl of the head homogenate supernatant for each sample was loaded into the plate. The plate was sealed and samples were incubated at 37°C for 90 min. Samples were then removed, the plate washed 4 times with 0.1% PBST. 100μl of a 1:2500 concentration of mouse anti-GFP antibody (Cell Signaling Life Technologies #A-11120) prepared in blocking buffer was added. The plate was sealed and left to shake overnight at 4°C. The plate was washed 4 times with 0.1% PBST and then 100μl of a 1:2000 concentration of donkey anti-mouse HRP (Jackson ImmunoResearch #715-035-150) in blocking buffer was added, the plate sealed and left to shake overnight at 4°C. The plate was washed 4 times with 0.1% PBST. 100μl of a 1:1 solution of TMB substrate and peroxide solutions (TMB substrate kit, Thermo #34021) added to each well. The plate was allowed to develop for 30 s – 2 min, depending on the time required for all samples to develop within the linear dynamic range of the assay. At this time, 100μl of 1N HCl was added to all wells. Optical density values at 450 and 630 nm were read on a colorimetric plate reader. OD values for each well were determined by subtracting the values from the 630nm wavelength from the 450nm reading. The average OD values for blank wells on the plate, in which PBST had been added in place of samples, was then subtracted from these values for each sample. These are the values reported as OD values throughout the paper. In some experiments, these were normalized to the average OD values of control samples, allowing comparison of results across plates. For additional details and about this protocol, see our previous methods paper that details this protocol and additional possible variations^24^.

### Screen

For the screen, we compiled a list of genes from existing literature and identified *Drosophila* orthologs using DiOPT. Lines to target these genes were identified from VDRC^48^ and up to two lines selected with < 2 predicted off-targets. These lines were crossed with flies expressing *Brp-GFP, repo-Gal4*. ≥ 9 male progeny of the desired genotype were collected at 3 dpe (48 to 72 hr after eclosion). 45 lines were deemed to be lethal, as they produced 1 or fewer progeny of the appropriate genotype and >100 flies eclosed. There were, in addition, several lines that we were unable to collect the total required 9 flies from, either because the proportion of the correct genotype produced was very low, or only females of the appropriate genotype were produced. These were not considered lethal and were excluded from our analyses.

Flies were frozen over time as they were collected (typically between ZT23 and ZT1) and stored at -80°C until use. For each screening sample, three fly heads were homogenized and then the lysates diluted 2x, such that the equivalent concentration of 1.5 fly heads was loaded. This was initially done to allow us to perform 2 technical replicates for each sample, but an interim analysis revealed that this conferred very little advantage in reducing sample variability, and therefore we used only one technical replicate per sample for the remaining screen. Three samples were prepared from flies from each genotype, representing 3 biological replicates. ELISAs and analysis were performed as described above, normalizing each sample to the average of the control samples (*brp-GFP, repo-Gal4* crossed with a VDRC isogenic control line – VDRC#60000) run on each plate. The fold change and p-value were determined using 1-way t-tests relative to 1. No corrections were made in p-values to correct for multiple comparisons, which with a cut-off of p<0.01 for ∼1000 lines could generate ∼10 false positive results. However, because we were quite limited in statistical power with an n of 3 per sample, we chose to be relatively liberal in these cut-off values, and opt to replicate potential hits in follow-up secondary screens. We categorized “hits” as those genes that when knocked down resulted in a >25% change from controls with a p<0.01 and “possible hits” that warrant further analysis as those with >15% change with p<0.05. Genes identified in the screen as “hits” or “possible hits” were subjected to gene ontology analysis using ShinyGo^92^. The top 30 differentially expressed pathways were analyzed and networks prepared from those categories that contained >3 genes in common.

### Imaging

For imaging, adult fly heads were removed from fly bodies and placed in a 4% PFA solution prepared in PBS for 20 min at room temperature. Heads were then briefly washed and dissected in PBS with 0.1% TritonX (0.1% PBST). Brains were kept in this solution on ice for up to 20 min. They were then transferred to 4% PFA prepared in PBS and fixed at room temperature for 20 min, and briefly washed in 0.1% PBST. For brains in which only endogenous fluorescent signal was to be evaluated, brains were then either transferred to Vectashield mounting media and mounted onto slides directly, or stored in 0.1% PBST at 4°C for up to 3 days, and then mounted. For brains that were subjected to staining, brains were instead transferred to 0.5 mL Eppendorf tubes and incubated in 0.5% PBST for 1 hour. Brains were then transferred into primary antibody solution (GAT 1:3000, developed by T. Stork), repo (8D12, developed by C. Goodman, 1:10), elaV (DSHB 7E8A10, 1:10), cleaved Dcp1 (Cell Signaling #9578, 1:1000), GFP (Life Technologies #A11120, 1:200) in 0.5% PBST at 4°C over 3 nights. Brains were then washed 3 times for 20^+^ min in 0.5% PBST and transferred to a secondary antibody solution (Alexa Fluors at 1:250 dilution) and incubated at 4°C over three nights. Brains were then washed 3 times, transferred to Vectashield mounting media and mounted onto slides.

Third instar larval and pupal brains collected at head eversion, ∼12 hr after puparium formation. Brains were dissected in PBS and transferred to 4% PFA on ice until all samples were collected. They were then transferred to RT and fixed for 20 additional min. Brains were then washed, transferred to Vectashield and mounted.

### LysoSensor

For LysoSensor experiments, heads were not prefixed, but brains dissected directly in live imaging buffer (Schneider’s Drosophila media + 1% pen/strep, 1% FBS and 10 μg/mL human insulin). A 1μM concentration of LysoSensor Blue (DND-167 #L7533) was added to the brains for 5min. Brains were then briefly washed two times with fresh live imaging buffer and mounted directly onto slides in this buffer. Imaging was performed within 30 min of dissection. Studies in which LysoSensor was co-localized with GAT were performed as described above, but after washing, brains were fixed in 4% PFA in PBS. These brains were then washed and subjected to immunostaining and imaging as described above.

### Imaging and analyses

All images were acquired on a Zeiss 880 confocal microscope, using traditional confocal imaging or Airyscan Fast imaging, as noted for each experiment below. For those image acquired with Airyscan, Airyscan processing was performed using automated settings in Zen Software. Image analyses were performed using Image J, except where noted below. Experimenters were blinded to genotype and time point when performing these analyses.

Mean intensity of Brp-GFP signal was evaluated across all the neuropil regions of the adult *Drosophila* brain that were accessible to imaging. Endogenous Brp-GFP signal was acquired using traditional confocal microscopy, with z sections taken every 5 μm through the anterior half of the brain, ∼10-12 slices. Below this depth, there was increasing difficulty in capturing Brp-GFP signal and thus we were unable to evaluate Brp-GFP levels in the most posterior neuropils. Because the neuropil structures are bilaterally symmetrical, Brp-GFP signal was quantified in one hemibrain for each fly. In each z plane acquired, each region of the neuropil was separately outlined and the mean intensity of Brp-GFP signal measured. All neuropils present in each z plane were quantified. Then the mean intensity values for each z slice were averaged to generate one mean intensity value for that region for each fly and region.

The percent area of FYVE-GFP and spn-RFP was quantified over time and when crq was knocked down from images acquired using traditional confocal microscopy. 5-7 z planes spaced 5 μm apart were acquired and max z projections performed. The entire brain area was then outlined as the ROI and the signal within manually thresholded. The percent positive area within this ROI was then measured. These are graphed for all groups as raw percentages in the figure and the same data were normalized to the 10-12 dpe and reported in the text. For analyses of FYVE-GFP and spn-RFP area in flies expressing UAS-crq, images were acquired using Airyscan Fast imaging. 5-7 z planes were acquired 5 μm apart. For analysis, 3 representative z slices were selected across these z slices. ROIs were drawn around brain and images manually thresholded. The percent positive area was quantified for each of these z slices individually and then averaged to generate the percent positive area measurement for each fly. These results were then normalized to the average value for 3 dpe controls for each genotype.

Engulfed synaptic material was analyzed by counting the number of mCherry^+^eGFP^-^ puncta present in brains from Brp^mCherry-eGFP^ flies. Images of the central brain were acquired using Airyscan Fast, with 7-9 z slices acquired 5 μm apart. eGFP and mCherry signals were overlayed and the mCherry^+^eGFP^-^ marked. These marks were cross-referenced against the mCherry and eGFP only channels to ensure that these puncta were only mCherry^+^. The number of mCherry^+^eGFP^-^ puncta across all acquired slices were quantified and then divided by the total area analyzed. Additional analyses were also performed to assess the eGFP+ area, the mCherry+ area and the ratio of mCherry+ to eGFP+ area across the central brain. For these analyses in adult fly brains, five z planes were evaluated, starting anteriorly where the antennal lobes were both fully visible, and spaced 5μm apart from that point. These values were then recorded for each z plane and then the average of these values calculated for each brain. For these analyses in larval and pupal brains, analyses were performed as above, except that the quantification was performed across 3 z planes across the VNC.

The number of Brp-GFP^+^ puncta were analyzed across three different experiments. In all experiments, images were acquired using Airyscan Fast imaging encompassing the region of interest with z sections taken 0.2 μm apart through this region. In flies expressing Brp-GFP endogenously across the brain, Brp-GFP puncta number was quantified in the antennal lobe. For each slice, this ROI was outlined and the area outside of the ROI cleared. We initially did this through the entire glomerulus, but pilot experiments demonstrated that analysis through a volume of 7-10 z slices gave very similar results to analysis through the entire volume, so only this 7-10 slice subset was used for analysis. The area of the ROI in each z section was measured and multiplied by 0.2 to obtain the volume of the area analyzed. To analyze puncta number, we scrolled to the middle of the z stack and adjusted the brightness and contrast to use the whole distribution of values and then converted the images from 16 to 8 bit. We then used the Auto Local Threshold function using the Bernsen method with a 12 px radius. After images were thresholded, we used the 3D objects counter analysis tool with a size filter of 10 – 2000, and selected to exclude objects on edges. Maps were shown so the experimenter could evaluate the accuracy of counting. The number on puncta was extracted from the results summary. This number was then normalized to the volume of the area analyzed.

For evaluating Brp-short-mCherry^+^ puncta number in the Or82a glomerulus, a very similar analysis protocol was used to that described above, except that it was not necessary to outline the region of interest because the synaptic label was already exclusively expressed in these neurons. We also analyzed synapse numbers across the entire glomerulus volume, to capture the total synapse number across the population, and therefore did not calculate or normalize to the volume of the ROI. This was similarly performed for Brp-short-GFP expressed in LN populations labeled by *GMR70A09-Gal4* and *GMR32F10-Gal4*.

For evaluating Brp-short-GFP^+^ puncta number in ppk neurons, we used Imaris analysis software. The “spot” function was used to automatically annotate GFP^+^ puncta across the volume of the image. This was then proofread by the experimenter and corrections made manually as needed. Imaris then reported the total number of spots across the population for each image. Like with the Or82a neurons, we did not normalize to volume of the ROI in this case, because synaptic labeling was restricted to ppk neurons.

Crq expression was analyzed from images in which *crq-Gal4* was used to drive the nuclear transcriptional reporter *UAS-H2B:mCherry*.*HA*. These images were acquired using traditional confocal imaging with 7-10 z planes acquired spaced 5 μm apart. Brains were co-stained with the glial nuclear marker repo as described above. mCherry and repo signals were overlayed. The number of all crq^+^, repo^+^crq^-^ and crq^+^repo^-^ nuclei were counted across 3 representative z planes. These numbers were then used to determine the percent of repo^+^ cells that were also crq^+^ and also the percent of *crq*^*+*^ cells that were repo^+^.

Neuronal death was evaluated by staining sections with elaV, a marker of neuronal nuclei, and cleaved Dcp1 to label dying cells. Images were acquired 2 μm apart through approximately half of the *Drosophila* brain to 50-70 μm in depth. Cleaved Dcp1^+^ cells were counted across this entire volume. Overlays of these two signals were used to determine whether Dcp1^+^ cells were neurons.

Neuronal nuclei were quantified across the central brain by counting the number of elaV+ cells across 5 selected z slices, starting with the first section in which both antennal lobes were distinguishable, and taken posteriorly every 4μm from that point. In each slice, the number of elaV+ nuclei were counted using the “analyze particles” function in Image J, using a range of 100-800 pixel units of size and a circularity of 0.5 – 1.0. The sum of all of these counts was obtained and this is the “number of elaV+ nuclei”.

LysoSensor signal was analyzed in images acquired using Airyscan Fast imaging. This experiment was performed in a background in which flies expressed *GMR25H07-Gal4, UAS-mCD8::GFP* which was used to allow clearer visualization of the neuropil. Three z planes 5 μm apart were imaged. Because this was done on live tissue rather than in mounting media, photobleaching prevented evaluation of this signal across a larger volume. LysoSensor^+^ area was evaluated within the neuropil of the antennal lobe, since pilot data suggested that this produced similar results to analysis across all the neuropil regions within our imaging window. The ROI outlining the antennal lobe was adjusted as needed to exclude imaging artifacts, in particular autofluorescent signal from the trachea. Thresholding to identify LysoSensor signal was performed manually and the percent area averaged across the 3 brain slices.

Brp-GFP area was performed in larvae and pupae. Images were acquired using Airyscan Fast and underwent Stiching before proceeding to Airyscan Processing using Zen Software. An ROI was drawn around the neuropil of the ventral nerve cord for each z plane and manual thresholding was used to capture the Brp-GFP^+^ area. This was done across 3 representative z planes and then averaged.

To evaluate clearance of Or85e neuronal debris after injury, an injury was performed in which one maxillary palp was removed from the fly. Seven days later, brains were dissected, fixed and stained for GFP to visualize the Or85e-mCD8::GFP signal. Images were acquired using traditional confocal imaging through the entire depth of the labeled glomeruli with z sections 1 μm apart. The volume of the glomerulus on the injured and uninjured size were determined by manually outlining the GFP^+^ area that marked each glomerulus through each z plane. These were then added together to determine the total volume for each glomerulus. The percent volume of the injured compared to the uninjured glomerulus was then calculated.

### Western blot

Flies were frozen and stored at -80°C until use. Heads were removed by vortexing and 3 heads per sample prepared in 1.5mL Eppendorf tubes. Samples were homogenized in PBS and Laemmeli buffer containing β-mercaptoethanol added. Samples were boiled for 5 min and spun down briefly. The supernatant was loaded into a 4-20% Tris-Bis Gel and run at 100V for ∼1 hr in Tris-glycine running buffer with 8% SDS. The gel was transferred to a PVDF membrane at 33V overnight at 4°C in a Tris-glycine transfer buffer containing 12% methanol. The membrane was blocked in 5% BSA in TBS with shaking for 1 hour at RT. The membrane was cut and incubated with nc82 antibody against Brp (DSHB nc82, 1:50), or β-tubulin antibody (DSHB E7, 1:1000) overnight at 4°C. Membranes were washed 3 times for 20 min in TBS and incubated in HRP conjugated secondary antibody (Jackson ImmunoResearch, 1:5000) in 5% BSA for 1 hour at RT with shaking. Membranes were washed 3 times for 20 min in TBS and excess buffer was then removed. ECL substrate (Super Signal West Pico Plus Chemiluminescent Substrate, Thermo #34577) was added to the membrane and blots imaged using a Chemidoc MP Imaging System. Optical density was quantified in Image J using the Gel Analyzer tool.

### Behavior

Within 24 hr of eclosion (0 dpe), flies were transferred to narrow vials containing cornmeal agar media. To blind scorers to genotype, vials were labeled with letter designations and genotypes contained in a separate key. Flies were separated by sex and 2-14 flies were housed per vial. While initially this number was 10-14, attrition to do death in some genotypes resulted in more variable conditions of housing density as the flies aged.Within 12 hr of behavioral testing, flies were transferred to a fresh vial.

To evaluate baseline climbing behavior, we marked each vial with a line 3 cm above the top of the food. Flies were tapped to the bottom of the vial and the number of flies which were above the 3 cm line at the reported intervals (5, 10, 15, 30 and 60s) were evaluated. The percentage of flies for each genotype and time point were calculated and plotted. > 100 flies per genotype were scored for each condition.

Seizure susceptibility was evaluated in flies by vortexing the vial at the max speed setting for 10 s. At the time points reported here, flies rarely exhibited complete sustained paralysis after vortexing, but we did see flies which appeared to exhibit recovery seizures, and, most commonly, a refractory period in which they remained at the bottom of the vial and failed to exhibit their typical anti-geotaxis behavior, scored as previously described^93^. All of these phenotypes were designated as “not recovered” while flies exhibiting normal anti-geotaxis were scored as “recovered”. These behaviors were scored 1 min after vortexing.

### Survival

Flies used for survival analysis were collected at 0 dpe and transferred to narrow vials containing cornmeal agar media. Flies were separated by sex and maintained < 14 flies per vial. While initially, there were 10-14 flies present per vial, at terminal time points for some genotypes, there were fewer flies. The number of remaining flies was counted every 2-3 days. Differences in survival curves were compared using log-rank Mantel-Cox tests.

### Statistics

All statistical analyses were performed using GraphPad Prism. All bar graphs are presented as mean +/-SEM and each dot represents one biological replicate (n). Outliers were identified using the Grubbs test with an alpha=0.05 and were excluded from further analyses. Normality was tested for each group in all datasets using a Shapiro-Wilk test with an alpha=0.05. Pairwise comparisons were performed using unpaired two-tailed t-tests when data were considered normal; Welch’s t-tests were used when data were not normally distributed. Ordinary one-way ANOVAs were performed to compare results from normal datasets across 3 or more groups. Additional post-hoc tests performed using Sidak’s, Tukey’s or Dunnett’s multiple comparisons tests, depending on what datasets were compared Datasets which were not normally distributed were compared using a Kruskal-Wallis test and Dunn’s post-hoc tests used. Groups for which post-hoc comparisons were made are designated on each graph as a bar between the two groups. Where no bar is present, these comparisons were not tested. For comparisons with two variables, two-way ANOVAs were performed and multiple comparisons made using Sidak’s post-hoc tests. p values are designated as ns, *p≥0.05, **p<0.01, *** p<0.001, and **** p<0.0001.

## Supplemental information titles and legends

**Figure S1: Developmental timecourse of additional synaptic proteins**

(A) Flies expressing an endogenously GFP tagged Syt7 (*syt7-GFP* **/ +**, males and females combined) were collected at the time points indicated. Three heads were pooled for each sample, Syt7-GFP OD values measured by ELISA, and then normalized to the average value of the 0 dpe group. Each n represents the measurement for one independent sample. The n for each group are: 19 (0 dpe), 20 (3 dpe), 20 (10-12 dpe). Data for all groups are normally distributed and compared using an ordinary one-way ANOVA (****) and Sidak’s post-hoc comparisons are indicated. (B) Flies expressing an endogenously GFP tagged dlg1 (*dlg1-GFP / + or dlg1-GFP / Y*, males and females normalized within sex and then combined) were collected at the time points indicated. Three heads were pooled for each sample, and GFP OD values measured by ELISA and were normalized to the average value of the 0 dpe group. Each n represents the measurement for one independent sample. The n for each group are: 18 (0 dpe), 19 (3 dpe), 20 (10-12 dpe). Data for all groups are normally distributed and were compared using an ordinary one-way ANOVA (****) and Sidak’s post-hoc comparisons are indicated.

**Figure S2: Glia engulf synaptic material**

(A) Synapses were labeled in flies expressing Brp endogenously tagged with an mCherry-eGFP dual fluorescent tag in controls (brp^mCherry-eGFP^ / +, GMR25H07-Gal4 / +, 10xUAS-FLP / +) and flies in which syntaxin 13 (syx13) is knocked down in astrocytes (astrocyte > syx13 RNAi - *brp*^*mCherry-eGFP*^ */ +, GMR25H07-Gal4 / +, 10xUAS-syx13 RNAi / +*). Male and female larvae were collected at wL3 and pupae collected at HE (∼12 hours after puparium formation). Ventral nerve cords were dissected and stained for eGFP and mCherry. Images were acquired with a 20x objective and (B) the percent eGFP and mCherry area was quantified across the entire area of the ventral nerve cord (VNC), from 3 different z planes. These results were then averaged to generate each data point, reflecting one VNC. eGFP area for each condition was normalized to the average value of the control larvae group. The n for each group are: 5 (control larvae), 5 (control puape), 6 (astro>syx13 RNAi larvae) and 6 (astro>syx13 RNAi pupae). Data from all groups was normally distributed, and a two-way ANOVA was performed (time point ****; genotype *** ; interaction **). (C) For each z plane, the ratio of mCherry : eGFP area was calculated, and then these results were averaged across the 3 z planes analyzed for each sample. These results were then normalized to the average value for the “control larvae” group. The n for each group are: 6 (control larvae), 6 (control puape), 6 (astro>syx13 RNAi larvae) and 6 (astro>syx13 RNAi pupae). Data from all groups was normally distributed and were compared using a two-way ANOVA (time point **; genotype **; interaction ***). (D) Additional analyses were performed on the images acquired from Fig 1I. The first 5 z planes were analyzed, starting where the antennal lobes were both fully visible and spaced 5μm apart. The percent area which was eGFP+ was recorded across the central brain, and averaged across these 5 z planes. Then these values were normalized to the average value from the 10-12 dpe group, and each n represents that value from one brain. The n for each group are: 9 (larvae), 10 (0 dpe), 10 (3 dpe), 10 (10-12 dpe). Data from all groups are normally distributed and analyzed using an ordinary one-way ANOVA (****) with Sidak’s post-hoc tests indicated. (E) The percent mCherry+ area was measured on the same samples and according to the same procedure as described for eGFP in D above. All groups were normally distributed and analyzed using an ordinary one-way ANOVA (****) with Sidak’s post-hoc tests indicated. (F) The ratio of mCherry+ area to eGFP+ area was calculated for each z plane from the data in D and E above and then averaged across all z planes. Data from all groups are normally distributed and compared using an ordinary one-way ANOVA (***) and Sidak’s post-hoc tests are indicated. (G) WT (VDRC isogenic control #60000) flies were collected at 3 dpe and dissected brains were incubated in LysoSensor Blue dye to identify acidic cellular compartments. Brains were then fixed and stained for the astrocyte membrane marker GAT. Images were acquired with a 63x objective and z planes were acquired 0.2μm apart across a total depth of 5-10μm. Astrocyte cell bodies are indicated with yellow asterisks. A white arrow is pointing to an example of LysoSensor+ signal within an astrocyte membrane and the gray arrow is pointing to an example of LysoSensor+ signal outside of an astrocyte membrane. Orthogonal projections of two example LysoSensor+ puncta identified within astrocyte membranes are shown. (H) An additional example of mCherry+eGFP-signal within an astrocyte membrane, as shown in Fig 1K.

**Figure S3: TRAPseq of astrocytes cross the lifespan**

(A) TRAPseq was performed on flies at the indicated time points. Ribosomes in astrocytes were labeled with eGFP (using male and female *alrm-Gal4/+, elav-Gal80/ +, UAS-L10a-eGFP/+* flies), GFP antibodies used to pull down these ribosomes and then the associated mRNA’s were sequenced. (C) Comparisons were made to identify differentially expressed genes between late pupal and 1-3 dpe astrocytes, (D) 1-3 vs 10-12 dpe astrocytes, and (E) 50-60 vs 10-12 dpe astrocytes.

**Figure S4: Validation of ELISA screening method to assay dynamic changes in Brp-GFP signal**

(A) Brp-GFP levels were measured from lysates from *brp/+* flies containing between ¼ and 10 fly heads and (B) signal could be readily detected in lysates even from ¼ of a fly head within the linear dynamic range of the assay (gray line – simple linear regression, R^2^ = 0.99; 2 independent samples are represented for each condition). For the rest of the assays in this figure, and for the screen, 3 fly heads were pooled for each sample. (C) ELISAs for Brp-GFP were performed with 20 replicates of control flies (*brp-GFP / +*) and the distribution of results compared between males and females. Male flies exhibited a less variable distribution. (D) 100 combinations of 1-5 samples were drawn from the 20 male control replicate samples and the distribution of these results plotted. Using one replicate resulted in the highest degree of deviation from the true sample mean, while five resulted in the least. We decided using three biological replicates appropriately balanced minimizing variability and required resource use. (E) Flies were collected at 10-12 dpe across the indicated zeitgeber times (ZT) and Brp-GFP levels measured across these samples by ELISA. This was performed in control flies (*brp-GFP / +*) or in *per*^*01*^ mutant flies crossed into the Brp-GFP line (*per*^*01*^ */ Y, brp-GFP / +*). Only males were used in these experiments. The number of samples for each group are: 13 (control, ZT2), 16 (control, ZT6), 14 (control, ZT14), 16 (control, ZT18), 4 (*per*^*01*^, *ZT2*), 4 (*per*^*01*^, ZT6), 5 (*per*^*01*^, ZT14), 5 (*per*^*01*^, ZT18). Data from two of these groups were not normally distributed (control, ZT14 and per^01^, ZT18). Therefore, we used a non-parametric Kruskal-Wallis test to analyze the effect of circadian time in control and *per*^*01*^ groups. (F) Circadian changes in Brp-GFP levels were evaluated in 3 dpe male brp-GFP / + flies across circadian times. The n for each group are: 4 (ZT2), 4 (ZT6), 4 (ZT14), 4 (ZT18). Data from all groups are normally distributed and were analyzed using an ordinary one-way ANOVA.

**Figure S5: Pathways enriched in genes required in glia for normal synapse development**

(A) Network analysis was performed on the top 30 differentially expressed pathways in the set of genes which, when knocked down in glia, increased Brp-GFP levels. The width of the edges representing the number of shared genes between these pathways. Roman numerals identify each cluster of related pathways, which (B) correspond to their location on the dendrogram below. (C, D) These analyses were repeated for genes that, when knocked down in glia, decreased Brp-GFP levels.

**Figure S6: Additional analyses of other screen hits**

(A) Ten additional genes from the screen were chosen for validation based on (1) involvement in a variety of different cellular processes, (2) being of potential interest because their orthologs have been implicated in human neurological disease and (3) representing a variety of screen outcomes (i.e. were initially very “strong” hits with low variance and high degree of difference from controls and “weak” hits which did not meet those criteria. *kug* did not meet criteria for a hit in the screen, but was identified as a possible gene of interest because its knockdown resulted in a 15% change in synapses with a p<0.05. These were first assessed for whether they similarly changed Brp levels when knocked down in glia using *repo-Gal4* in female flies (as the initial screen was performed in males), and using additional glia-specific drivers (*GMR25H07-Gal4* which is a strong astrocyte driver and *alrm-Gal4* which is a weaker astrocyte driver). For each condition, 3 heads were pooled for each sample and OD values for GFP measured by ELISA for each sample. An n=3 was run for each gene and condition. These were normalized to the average value of the controls run for each group, where the drivers were crossed with the VDRC isogenic control line #60000 (for 3 dpe female all glia *brp-GFP / +, repo-Gal4 / +*; for 3 dpe male strong astrocytes *brp-GFP / +, GMR25H07-Gal4 / +;* for 3 dpe male weak astrocytes *brp-GFP / +, alrm-Gal4 / +*). The mean values were averaged and are reported here using a heat map. No change from control is represented as black (1.0), an increase in GFP levels up to 1.5-fold shown in blue and decreases in GFP levels down to 0.5-fold compared to controls shown in pink. Knockdown of *Fkbp59, btl* and *CG14886* all increased Brp values in the initial screen, and also showed increases across these other conditions. *Acox57D-d* and *CAP* knockdown resulted in decreases in Brp-GFP levels in the initial screen and showed decreases, albeit to smaller degrees in some conditions, in these experiments. Interestingly knockdown of *kul, Loxl1* and *Ser7* all increased Brp-GFP levels in the screen, and when knocked down in all glia in females, but showed the opposite effect on Brp-GFP levels when knocked down solely in astrocytes. *Col4a1* and *kug* did not produce consistent changes in Brp-GFP levels using these additional measures. (B) Imaging was performed using a 40x objective with z slices taken from the anterior-most part of the brain and z sections were acquired 5μm through 60μm total depth. The entire area of the central brain (outlined in white) was imaged. In each z section, either (C) the entire central brain or (D) an antennal lobe was defined as the ROI and the mean intensity of GFP signal within this ROI determined. These values were averaged across all z slices for each brain. This number was then normalized for each knockdown condition (*brp-GFP / +, repo-Gal4 / +, UAS-RNAi / +* (against the genes indicated on the x axis) to the average value for the control (*brp-GFP / +, repo-Gal4 / +*) group. Each n represents one brain. The n for each group are: 7 (control), 7 (Fkbp59), 4 (btl), 3 (CG14866), 4 (Acox57D-d), 5 (CAP), 4 (kul), 3 (Loxl1), 8 (Ser7), 5 (Col4a1), 5 (kug). All groups were normally distributed. Ordinary one-way ANOVAs were performed (**** for central brain and **** for antennal lobes) and each group was compared to the control group using a Dunnett’s multiple comparisons test, indicated in red.

**Figure S7:**

(A) A diagram of the *crq* locus with the regions targeted by the RNAi lines used in this study, and the *crq* mutant lines are shown. (B) A western blot was performed to assess Brp protein levels in control (*yw*), crq^CR02509^ (*crq*^*CR02509*^) and crq^CR02509^, crq>UAS-crq (*crq*^*CR02509*^, *UAS-crq / +*). Five heads were pooled for each sample and male and female flies at 3 dpe were used in this experiment. (C) Brp signal was measured and normalized to β-tubulin for each sample. This was then normalized to the average value of the control group. An n of 3 samples was evaluated for each group. All groups were normally distributed and compared using an ordinary one-way ANOVA (*) with results from Sidak’s posthoc tests shown. (D) Male and female flies were collected at 3 dpe for control (*brp-GFP / +, repo-Gal4 / +*), crq RNAi#1 (*brp-GFP / +, repo-Gal4 / +, UAS-crq RNAi#1 / +*) and crq RNAi#2 (*brp-GFP / +, repo-Gal4 / +, UAS-crq RNAi#2/ +*) groups. Images were acquired using a 20x objective and the entire central brain and one optic lobe were imaged for each brain. z sections were acquired 2μm apart from the anterior-most portion of the brain to a depth of ∼40μm. z projections were prepared and each of the anterior neuropil regions outlined (labeled with roman numeral in diagram and corresponding to the roman numerals shown below the x-axis on the graph). Within each neuropil region, the area was outlined and the mean gray value determined. Because the neuropils are bilaterally symmetrical, we obtained two values for each brain region for the central brain neuropil regions and these were averaged together to generate a value for each brain. For analysis of the optic lobe (V), only one neuropil was analyzed per brain. The n for each group are: 16 (control), 10 (crq RNAi#1), 14 (crq RNAi#2). All groups were normally distributed except for the control group for neuropil region VI. Ordinary one-way ANOVAs were performed across groups for each neuropil regions I-V. These results were: I (**), II (***), III (**), IV (**), V (*). Results from Sidak’s post-hoc tests are indicated. A Kruskal-Wallis test was performed across groups for neuropil region VI (ns). Dunn’s post-hoc tests are shown.

**Figure S8:**

(A) A *crq-Gal4* line was used as a transcriptional reporter to drive a nuclear-localized mCherry (*crq-Gal4 / +, UAS-H2B:mCherry*.*HA / +*, magenta), and glial nuclei were stained with repo (green) in male and female flies. Images were acquired using a 40x objective and images acquired every 2μm across the entire central brain. Representative images are single z slices. In the merged insets, most nuclei are double positive, but a few are mCherry+ only (indicated by the black arrow) or repo+ only (indicated by the white arrow). (B) The number of *crq*+ cells, repo+*crq*-cells and *crq*+repo-cells were counted across 3 representative z sections 4μm apart. The number of cells across these 3 sections were added, and then this was used to calculate the percent *crq* expressing cells which are glia (*crq*+repo+ cells / total *crq*+ cells) and (C) the percent of glia expressing *crq* (*crq*+repo+ cells / total repo+ cells) were quantified at the ages indicated. Each n represents one individual fly brain. The n for each group are: 9 (0 dpe), 7 (3 dpe) and 4 (10-12 dpe). Data from all groups was normally distributed. Ordinary one-way ANOVAs were used to compare the groups in (B - ns) and (C ****) and Sidak’s post-hoc tests are indicated. (D) Flies expressing Brp-GFP were collected from male and female flies at 3 dpe. Three heads were pooled for each sample and GFP levels in each sample were measured by ELISA. Crq was knocked down in all glia (*repo-Gal4*), neurons (*elaV-Gal4*), astrocytes (*alrm-Gal4*), ensheathing glia (*GMR56F03-Gal4*) and cortex glia (*cortex glia split-Gal4*) and Brp-GFP levels measured and normalized to a control in which the indicated Gal4 was expressed alone (dotted line). The n for each group are: 9 (all glia), 9 (astrocytes), 4 (neurons), 5 (ensheathing glia), and 5 (cortex glia). The data for all groups are normally distributed. A one-sample t test was performed for each group relative to the hypothetical value of 1.

**Figure S9:**

(A) Brains of male and female flies at the indicated ages (control – *repo-Gal4 / +* and crq RNAi#1 – *repo-Gal4 / +, UAS-crq RNAi#1 / +*) were stained to visualize neuronal nuclei (elaV, green) and the nuclei of dying cells (cleaved Dcp1, magenta). The zoomed out images are z projections through a portion of the central brain, and the insets below are single z planes. These images were acquired at 40x from the anterior-most portion of the central brain to a depth of ∼60μm, with z planes spaced 5μm apart. (B) All z planes were evaluated for the presence of cleaved Dcp1+ cells. There were no cleaved Dcp1+elaV+ cells identified, but the total number of cleaved Dcp1+ cells was quantified. Each n represents one brain. The n for each group are: 12 (control, 3 dpe), 9 (crq RNAi#1, 3 dpe), 20 (control, 10-12 dpe), 20 (crq RNAi#1, 10-12 dpe). (C) Brains of male and female flies were collected at the indicated time points and stained for elaV (green) to identify neuronal nuclei. Images of the whole area of the central brain were imaged with a 40x objective with z sections 2μm apart to a depth of ∼60μm. This imaging was performed in control (*repo-Gal4 / +*) and glia>crq RNAi#1 (*repo-Gal4 / +, UAS-crq RNAi#1 / +*) groups; (D) control (*GMR25H07-Gal4 / +*) and astro>crq RNAi#1 (astrocytes; *GMR25H07-Gal4 / +, UAS-crq RNAi#1 / +*) groups; and (E) control (*Ctx-glia-split-Gal4 / +*) and cortex glia>crq RNAi#1 (*Ctx-glia-split-Gal4 / +, UAS-crq RNAi#1 / +*) groups. (F) These images were analyzed to evaluate neuron number by selecting 5 z slices, starting with the first section in which both antennal lobes were distinguishable, and taken posteriorly every 4μm from that point. In each slice, the number of elaV+ nuclei were counted. The sum of all of these counts was obtained and this is the “number of elaV+ nuclei” graphed. Each n represents one brain. The n for each group are: 11 (control, 3 dpe), 13 (control, 10-12 dpe), 11 (glia>crq RNAi#1, 3 dpe), 15 (glia>crq RNAi#1, 10-12 dpe). Data from all groups are normally distributed. Groups were compared using a two-way ANOVA (age ****, genotype – ns, interaction, ns) and results from Sidak’s post-hoc tests are indicated. (G) Images were analyzed as described in (F). The n for each group are: 9 (control, 3 dpe), 11 (control, 10-12 dpe), 10 (astro>crq RNAi#1, 3 dpe), 11 (astro>crq RNAi#1, 10-12 dpe). Data from all groups are normally distributed. Groups were compared using a two-way ANOVA (age ****, genotype **, interaction – ns) and Sidak’s post-hoc tests are shown. (H) Images were analyzed as described in (F). The n for each group are: 12 (control, 3 dpe), 12 (control, 10-12 dpe), 10 (cortex glia>crqRNAi#1, 3 dpe), 9 (cortex glia>crqRNAi#1, 10-12 dpe). Groups are normally distributed except for the 3 dpe cortex glia>crqRNAi#1 group. Groups were compared using a two-way ANOVA (age ****, genotype – ns, interaction – ns). Sidak’s post-hoc tests are shown.

**Figure S10:**

(A) LysoSensor was used to visualize lysosomes in acutely dissected live fly brains at 3 dpe; while there was some background from tracheal signal which was excluded from the analysis, individual puncta representing lysosomal signal, as shown in the zoomed in insets below, was quantified within the neuropil. Representative images shown here are from a single z plane. Additional details are provided in Figure 1D. (B) Astrocyte membranes were labeled using the *GMR25H07-Gal4* driver to express *UAS-mCD8::GFP* and representative images from antennal lobes are shown. This was performed in male and female flies at 3 dpe from control (*GMR25H07-Gal4 / +, UAS-mCD8::GFP / +*), astro> crq RNAi#1 (*GMR25H07-Gal4 / +, UAS-mCD8::GFP / +, UAS-crq RNAi#1 / +*) and astro> crq RNAi#2 (*GMR25H07-Gal4 / +, UAS-mCD8::GFP / +, UAS-crq RNAi#2 / +*) groups. Images were acquired with a 63x objective in the antennal lobes starting at the anterior-most portion of the brain and z sections were acquired every 3μm apart to a total depth of 24μm. Representative images are z projected through the entire z stack. (C) GFP signal from these genotypes was quantified by ELISA. Three brains were pooled for each sample from male and female flies at 3 dpe. The n for each group are: 8 (control), 8 (crq RNAi#1) and 7 (crq RNAi#2). Data from all groups are normally distributed. Groups were compared using an ordinary one-way ANOVA (ns) and Sidak’s post-hoc tests are shown. (D) Additional analysis measures were performed on the images shown in Figure 4H, in control (*brp*^*mCherry-eGFP*^ / +, *repo-Gal4 / +*), glia> crq RNAi#1 (*brp*^*mCherry-* eGFP^ / +, *repo-Gal4 / +, UAS-crq RNAi#1 / +*) and glia> crq RNAi#2 (*brp*^*mCherry-eGFP*^ / +, *repo-Gal4 / +, UAS-crq RNAi#2 / +*) groups. The percent eGFP area was evaluated in the central brain from 5 z sections across the central brain. This was then averaged across these 5 images for each brain. Each n represents one brain. The n for each group are: 6 (control), 7 (crq RNAi #1), 3 (crq RNAi #2). Data from all groups are normally distributed and compared using an ordinary one-way ANOVA (*) and Dunnett’s post-hoc tests are shown. (E) mCherry+ area was quantified from the same set of images as described above. The n for each group are: 6 (control), 7 (crq RNAi#1), 4 (crq RNAi#2). All data are normally distributed and compared using an ordinary one-way ANOVA (ns) and Dunnett’s post-hoc tests are shown. (F) The ratio of mCherry to eGFP area was calculated for each z section and then this was averaged across all 5 sections analyzed. The n for each group are: 6 (control), 7 (crq RNAi #1) and 4 (crq RNAi#2). All data are normally distributed and compared using an ordinary one-way ANOVA (****) and results from Dunnett’s post-hoc tests are shown.

**Figure S11:**

(A) Climbing behavior was quantified in male and female *yw* control and *crq*^*MI12323*^ mutants and (B) control (*repo-Gal4 / +*) flies and flies in which *crq* was knocked down in glia (*repo-Gal4 / +, UAS-crq RNAi / +*). The percentage of flies that had climbed > 3cm in the time indicated was quantified in >100 flies for each genotype and time point. (C) Climbing behavior was similarly recorded in male and female control (*repo-Gal4* / +) and crq RNAi#1 (*repo-Gal4 / +, UAS-cra RNAi#1 / +*) flies which were raised at 29°C starting within 1 day of eclosion. (D) Survival was recorded in *crq* mutant and *yw* controls and (E) male only control (*repo-Gal4 / +*) flies and flies in which *crq* was knocked down in glia (*repo-Gal4 / +, UAS-crq RNAi / +*), in either a *w*^*1118*^ or *Sh*^*5*^ mutant background. Chi square tests were used to make comparisons between groups and results are indicated.

**Figure S12:**

(A) Brp-GFP levels were measured by ELISA in aged (30-35 dpe) male and female flies. Two heads were pooled for each sample and GFP levels measured in each sample from the following groups: control (*brp-GFP / +, repo-Gal4 / +)*, crq RNAi#1 (*brp-GFP / +, repo-Gal4 / +, UAS-crq RNAi#1 / +*) and crq RNAi#2 (*brp-GFP / +, repo-Gal4 / +, UAS-crq RNAi#2 / +*). OD values for each group were normalized to the average value of 10-12 dpe controls. The n for each group is 10 samples. All data are normally distributed and were compared using an ordinary one-way ANOVA (**). Results from Dunnett’s post-hoc tests are shown. (B) Brp-GFP levels were measured by ELISA in 30-35 dpe male and female flies as described above to test the effect of knocking down *crq* in astrocytes in the following genotypes: control (*brp-GFP / +, GMR25H07-Gal4 / +)*, crq RNAi#1 (*brp-GFP / +, GMR25H07-Gal4 / +, UAS-crq RNAi#1 / +*) and crq RNAi#2 (*brp-GFP / +, GMR25H07-Gal4, UAS-crq RNAi#2 / +*). OD values for each group were normalized to the average value of 10-12 dpe controls. The n for each group are: 14 (control), 7 (crq RNAi#1) and 20 (crq RNAi#2). Data from all groups are normally distributed. Groups were compared using an ordinary one-way ANOVA (****) and results from Dunnett’s post-hoc tests are shown. (C) Brp-GFP levels were measured by ELISA to evaluate synaptic protein changes with age when knocking down *crq* in cortex glia. Male and female flies were collected at 30-35 dpe and ELISAs performed as described above. The genotypes used in this experiment are: control (*brp-GFP / +, Ctx-glia-split-Gal4 / +)*, crq RNAi#1 (*brp-GFP / +, Ctx-glia-split-Gal4 / +, UAS-crq RNAi#1 / +*) and crq RNAi#2 (*brp-GFP / +, Ctx-glia-split-Gal4 / +, UAS-crq RNAi#2 / +*). OD values for each group were normalized to the average value of 10-12 dpe controls. The n for each group are: 16 (control), 10 (crq RNAi#1), 13 (crq RNAi#2). Data from all groups are normally distributed. Groups were compared using an ordinary one-way ANOVA (***) and results from Dunnett’s post-hoc tests are shown. (D) 30-35 dpe brains were collected and stained for cleaved DCP1 (magenta) to identify apoptotic cells and elaV (green) to identify neuronal nuclei. Male and female brains were used. Images were acquired using a 40x objective from the anterior-most portion of the central brain to a depth of ∼60μm, with 5μm z steps. The images shown are from single z planes and show examples of cleaved DCP1+elaV+ cells (white arrow) and cleaved DCP1+elaV-cells (gray arrow). Cleaved DCP1 signal was also evident in the neuropil of several aged brains. Some examples appeared punctate and diffuse across the neuropil, while in others cleaved DCP1 seemed to label neurites in varying stages of degeneration. (E) All z planes were evaluated for the presence of cleaved Dcp1+ nuclei. The total number across all z planes was quantified for each brain. The n for each group are: 14 (control), 14 (crq RNAi#1), 15 (crq RNAi#2). Data from all groups are normally distributed. Groups were compared using an ordinary one-way ANOVA (ns) and Dunnett’s post-hoc tests are shown. (F) Analyses were performed as in E for cells which were cleaved DCP1+ and elaV+. The n for each group are: 14 (control), 14 (crq RNAi#1), 15 (crq RNAi#2). All groups are normally distributed and compared using an ordinary one-way ANOVA (ns) and Dunnett’s post-hoc tests are indicated. (G) Brains of male and female flies were collected at the indicated time points and stained for elaV to identify neuronal nuclei. Images of the whole area of the central brain were imaged with a 40x objective with z sections 2μm apart to a depth of ∼60μm. This imaging was performed in control (*repo-Gal4 / +* at both 10-12 and 30-35 dpe), crq RNAi#1 (*repo-Gal4 / +, UAS-crq RNAi#1 / +*) and crq RNAi#2 (*repo-Gal4 / +, UAS-crq RNAi#2 / +*) groups. (E) These images were analyzed to evaluate neuron number by selecting 5 z slices, starting with the first section in which both antennal lobes were distinguishable, and taken posteriorly every 4μm from that point. In each slice, the number of elaV+ nuclei was counted. The sum of all of these counts was obtained and graphed. Each n represents one brain. The n for each group are: 13 (10-12 dpe) and 11 (30-35 dpe). Both groups are normally distributed. Data were compared using an unpaired t-test. (I) The number of nuclei for each group (crq RNAi#1, 30-35 dpe and crq RNAi#2, 30-35 dpe) was plotted and the blue dotted line shows the average value from the control 30-35 dpe group from H. The n for each group are: 13 (crq RNAi#1) and 14 (crq RNAi#2). Data from both groups are normally distributed. Each group was analyzed using a one-sample t-test relative to the hypothetical mean of the control 30-35 dpe group average.

**Figure S13:**

(A) Brp-GFP expressing flies (control - *brp-GFP / +, GMR25H07-Gal4 / +, 10xUAS-FLP / +* and crq RNAi#1 – *brp-GFP / +, GMR25H07-Gal4 / +, UAS-crq RNAi#1 / +*) were dissected at larval (wL3) and pupal (head eversion, ∼12hr after puparium formation) stages. Animals of both sexes were evaluated. Images were acquired using a 20x objective and (B) the percent Brp-GFP^+^ area quantified across 3 z planes, and these results averaged for each brain. Each n represents one brain. The n for each group are: 2 (control, Larva), 4 (control, Pupa), 5 (crq RNAi#1, Larva), 5 (crq RNAi#1, Pupa). Where possible to test, groups are normally distributed. Groups were compared using a two-way ANOVA (age ****, genotype – ns, interaction – ns). (C) Maxillary palp injuries were performed unilaterally in male and female flies and fly brains were dissected and fixed 7 days after injury. Or85e^+^ neurons which project to the imaged glomerulus were genetically labeled and stained with GFP. Images were acquired at 40x and z planes acquired every 1μm through the depth of the labeled glomerulus. z projections are shown in representative images. (D) The volume of each glomerulus was determined by outlining the GFP+ region on each side of the brain for each z section. The percent volume of the injured glomerulus compared to the uninjured glomerulus was then calculated. Each brain represents one n. The n for each group are: 24 (control) and 16 (crq RNAi#1). Data are normally distributed. An unpaired t-test was used to compare these groups. (E) ELISAs were used to evaluate Brp-GFP levels in flies in control flies (*brp-GFP / +, repo-Gal4 / +*) and flies in which *drpr* was knocked down in glia (*brp-GFP / +, repo-Gal4 / +, UAS-drpr RNAi / +*). Samples were prepared from male and female flies at the time points indicated. Two heads were pooled for each sample and GFP levels measured by ELISA. OD values were then normalized to the average value for the 10-12 dpe group. The n for each group are: 7 (10-12 dpe) and 6 (30-35 dpe). Data are normally distributed and were compared using an unpaired t-test. (F) Transcripts per million (TPM) reads were analyzed for *crq* and *drpr* from an RNA sequencing dataset from Purice et al., 2017. The n for each group is 5. All groups are normally distributed. Unpaired t-tests were performed between “No injury” and “Injury” groups for *crq*, and also for “No injury” and “Injury” groups for *drpr*. Results from those comparisons are indicated. (G) Read per Million (RPM) reads were analyzed for *crq* and *drpr* from an RNA sequencing dataset from Pacifico et al., 2018. The n for each group is 6, except for the drpr 40 dpe group which has an n of 5 due to a significant outlier in the dataset. Data from all groups was normally distributed. Unpaired t-tests were performed between 5 dpe and 40 dpe groups for *crq* and also for *drpr*.

**Table S1: Astrocyte TRAPseq data**

**Table S2: Screen data**

