## Supplemental Figures (S1 - S13) for "Developmental and age-related synapse elimination is mediated by glial Croquemort"

Figure S1: Developmental timecourse of additional synaptic proteins

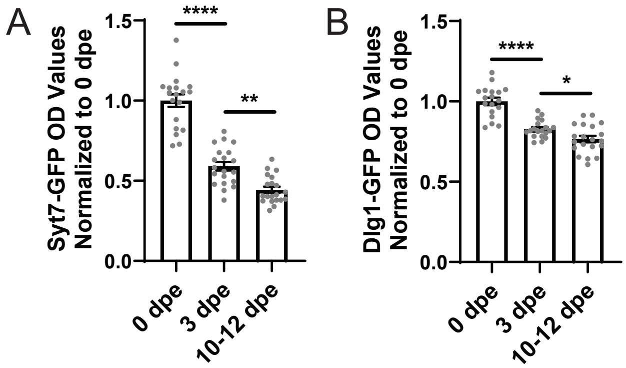

Figure S2: Glia engulf synaptic material

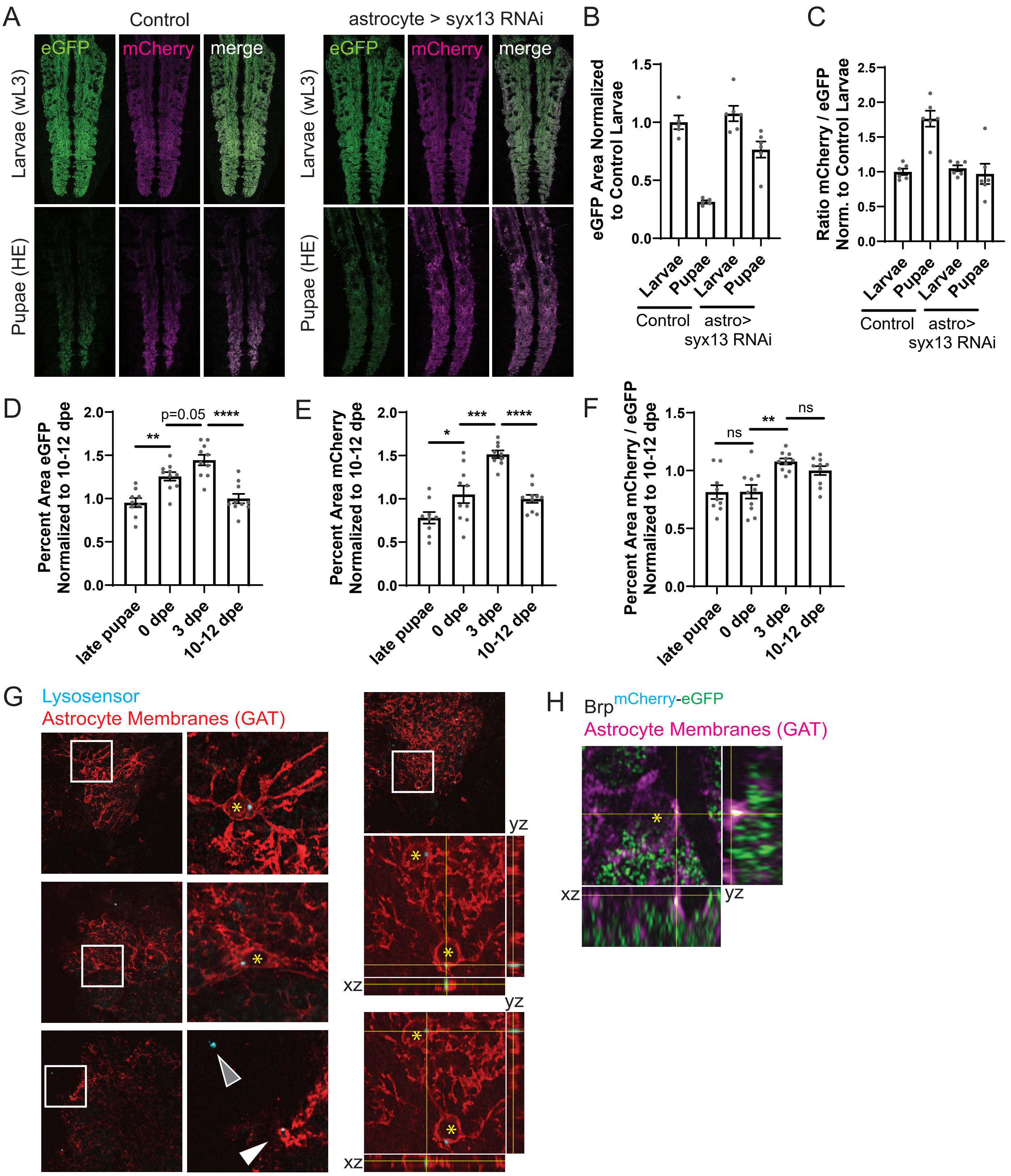

Figure S3: TRAPseq of astrocytes across the lifespan

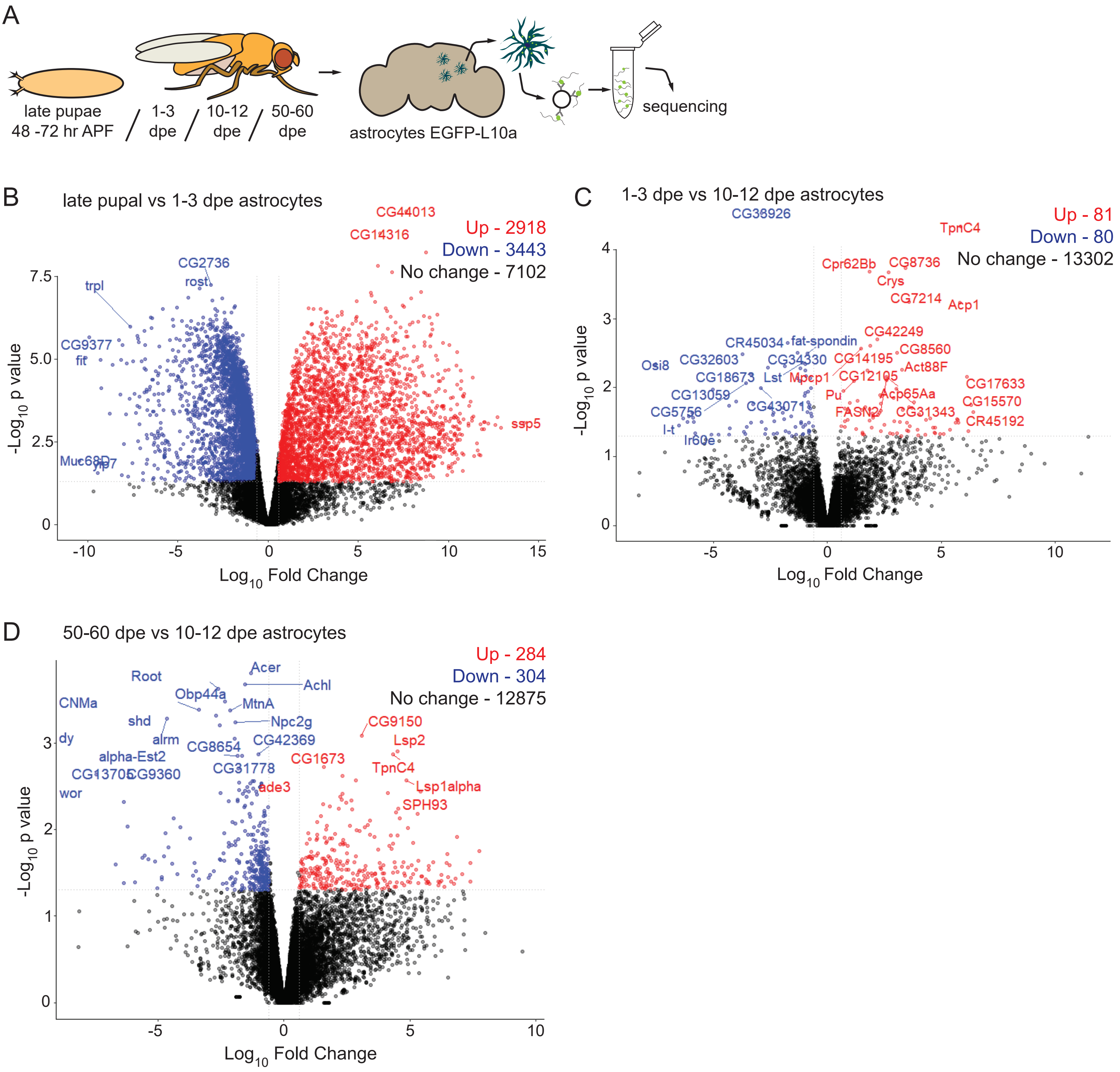

Figure S4: Validation of ELISA screening method to assay dynamic changes in Brp-GFP signal

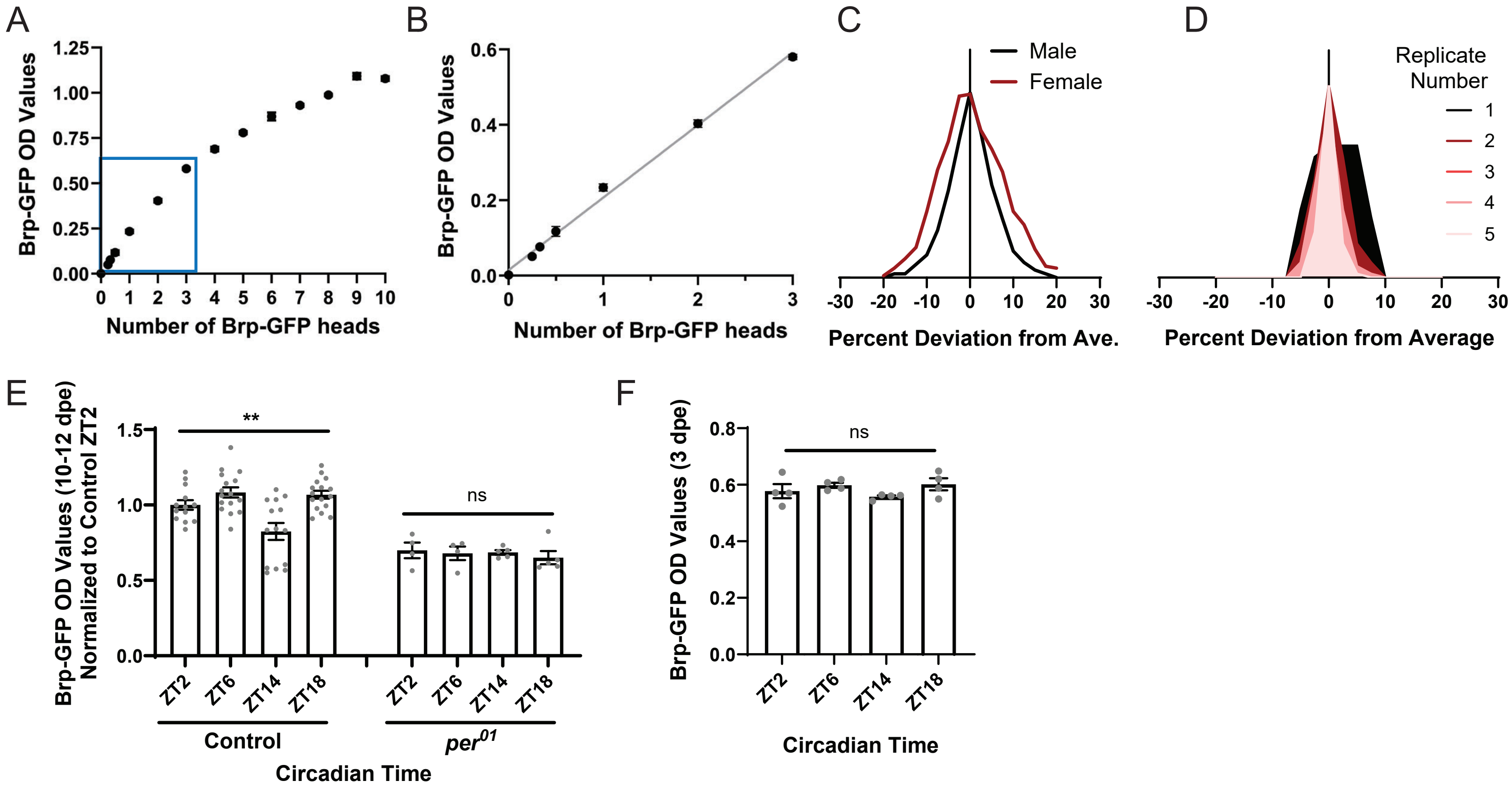

Figure S5: Pathways enriched in genes required in glia for normal synapse development

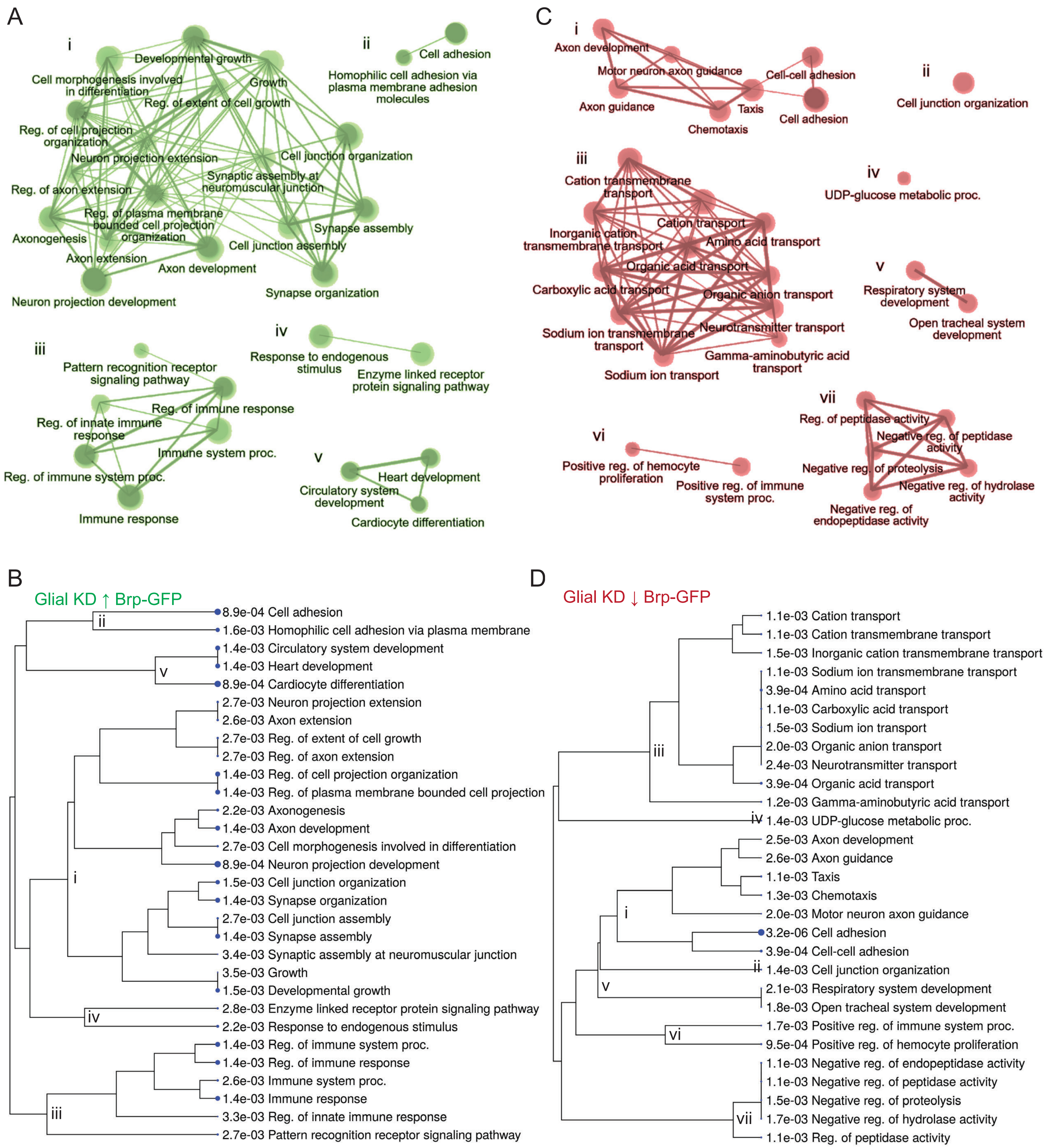

Figure S6: Additional analyses of other screen hits

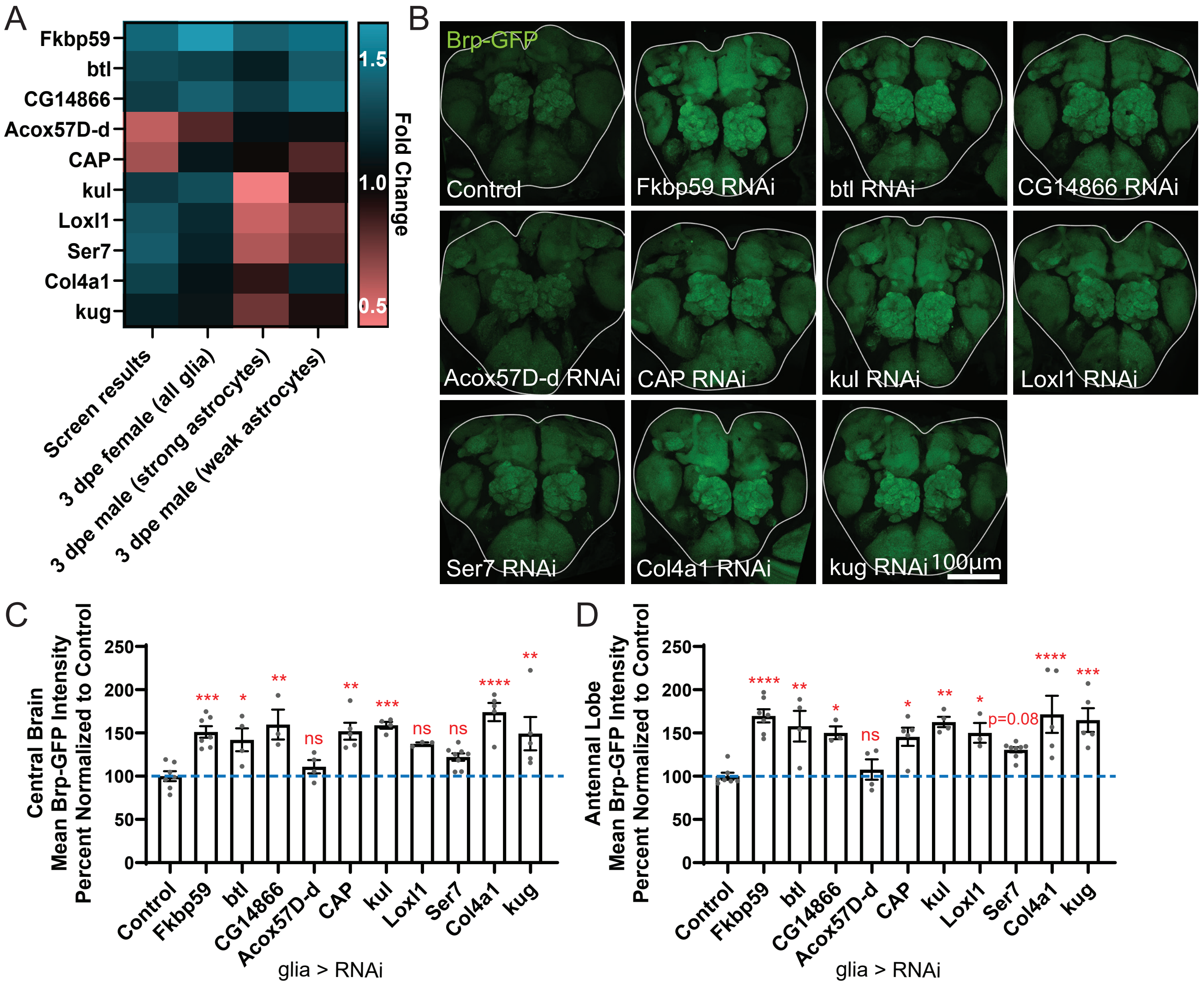

Figure S7: Crq is required in glia for normal synaptic protein levels

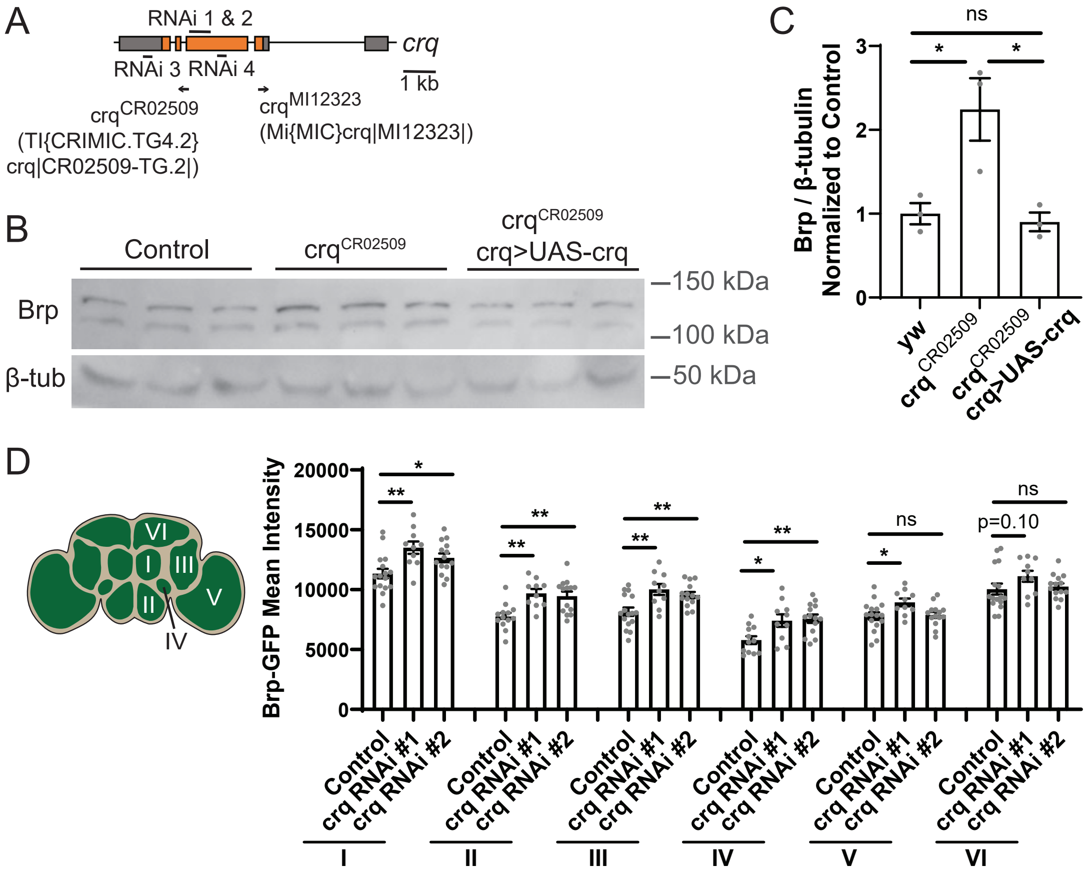

Figure S8: Crq expression over development

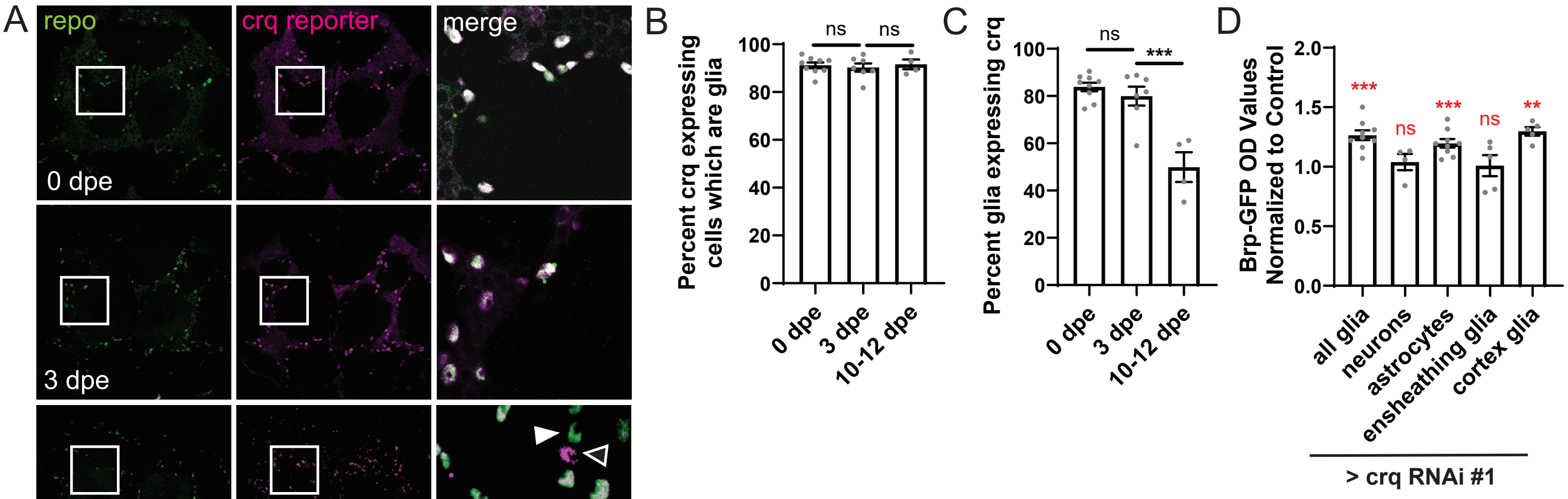

Figure S9: Crq is not required for developmental decreases in neuron numbers

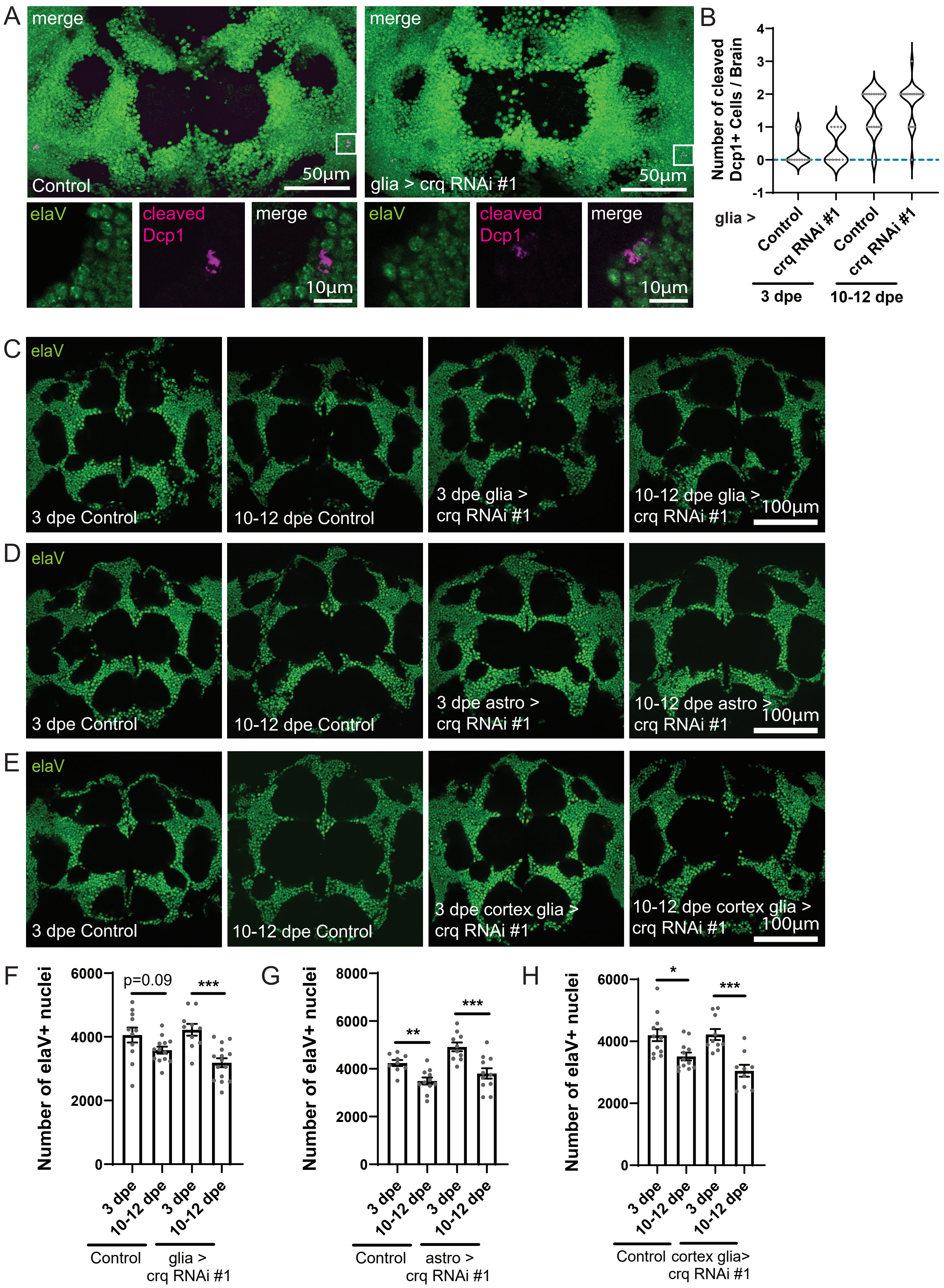

Figure S10: Loss of crq attenuates phagocytic markers in glia

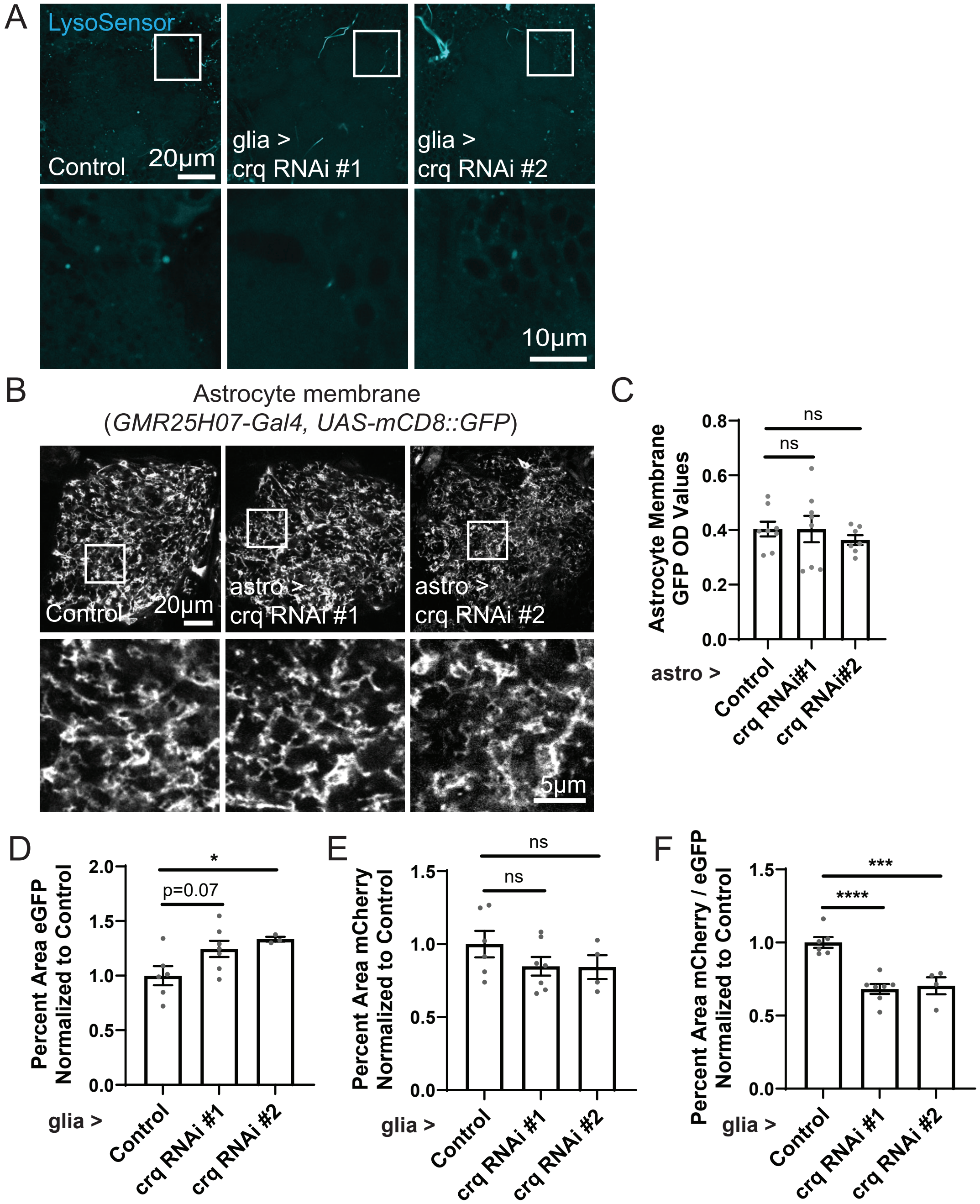

Figure S11: Climbing behavior and survival are altered with loss of Crq

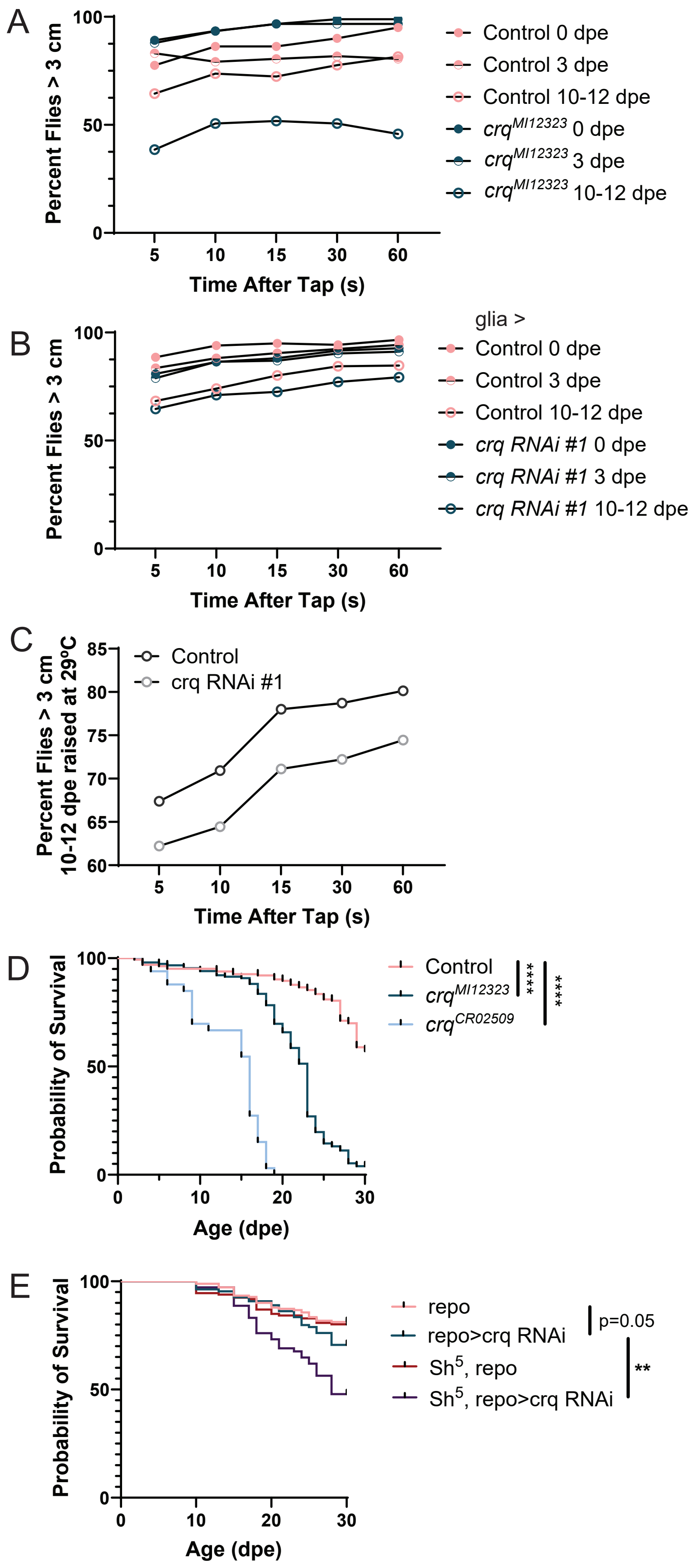

Figure S12: Glial Crq is required for synapse, but not neuronal loss with age

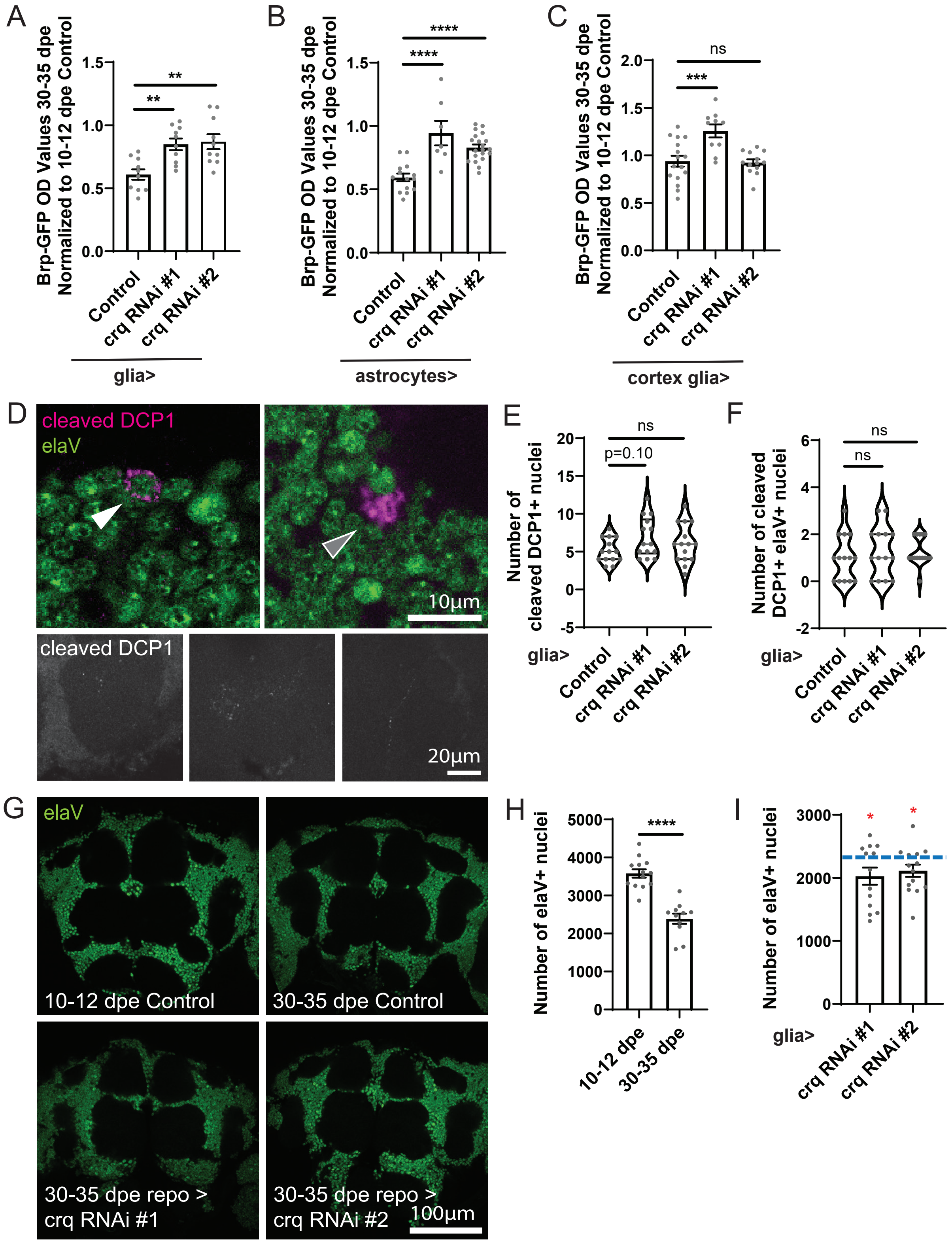

Figure S13: Crq is not required for glial engulfment during metamorphosis or after injury

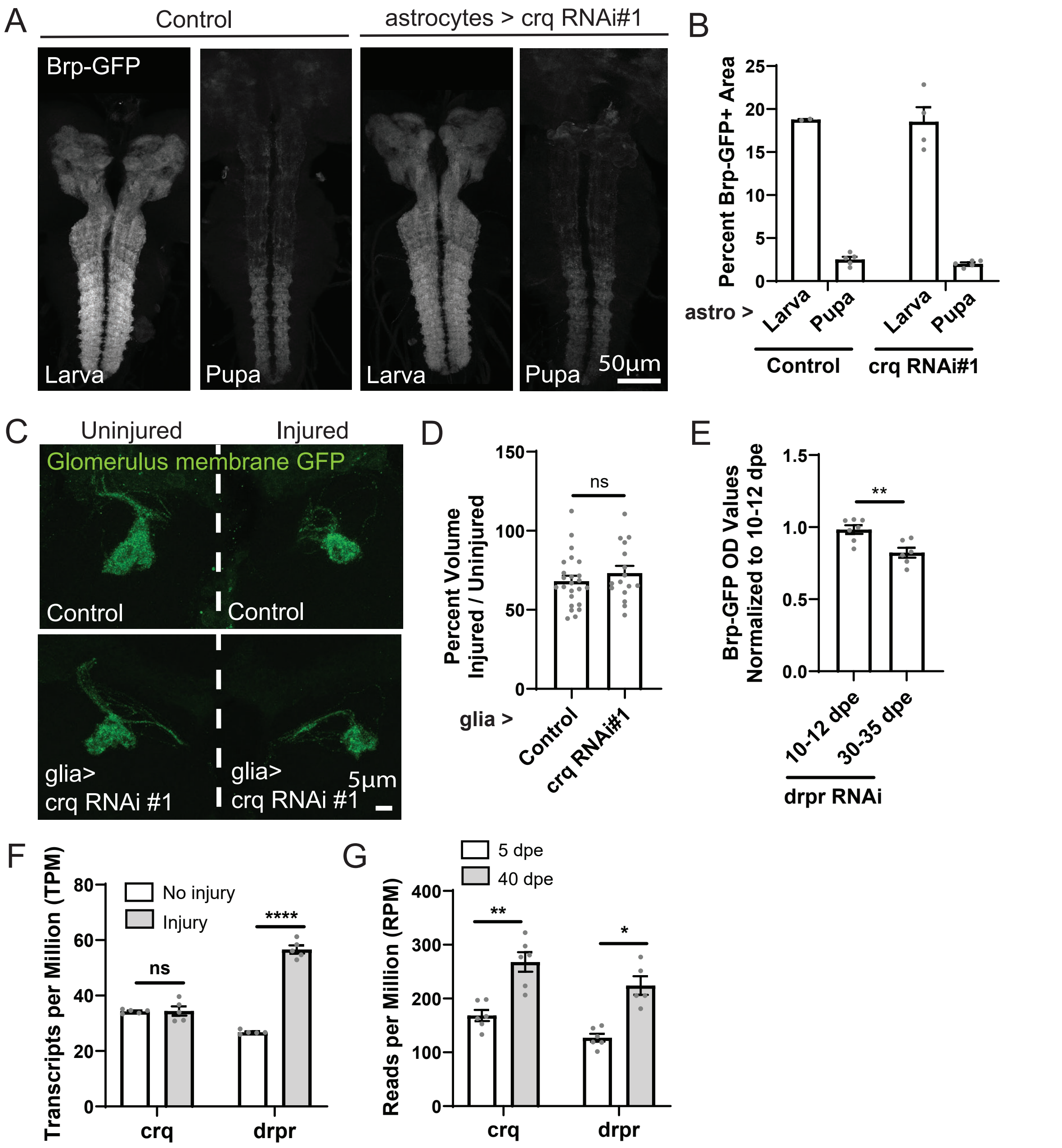
